# Two new immature and dysfunctional neutrophil cell subsets define a predictive signature of sepsis useable in clinical practice

**DOI:** 10.1101/2020.05.29.123992

**Authors:** Aïda Meghraoui-Kheddar, Benjamin G. Chousterman, Noëlline Guillou, Sierra M. Barone, Samuel Granjeaud, Helene Vallet, Aurélien Corneau, Karim Guessous, Alexandre Boissonnas, Jonathan M. Irish, Christophe Combadière

## Abstract

Sepsis is the leading cause of death in adult intensive care units. At present, sepsis diagnosis relies on non-specific clinical features. It could transform clinical care to have objective immune cell biomarkers that could predict sepsis diagnosis and guide treatment. For decades, neutrophil phenotypes have been studied in sepsis, but a diagnostic cell subset has yet to be identified. Here, high dimensional mass cytometry was used to reveal for the first time a specific neutrophil signature of sepsis severity that does not overlap with other inflammatory biomarkers, and that distinguishes patients with sepsis from those with non-infectious inflammatory syndrome. Unsupervised analysis of 42-dimesional mass cytometry data characterized previously unappreciated heterogeneity within the CD64^+^ immature neutrophils and revealed two new subsets distinguished by CD123 and PD-L1 expression. These immature neutrophils exhibited diminished activation and phagocytosis functions. The proportion of CD123-expressing neutrophils also correlated with clinical severity. Critically, this study showed that these two new neutrophil subsets were specific to sepsis and detectable by routine flow cytometry using seven markers. The demonstration here that a simple blood test distinguishes sepsis from other inflammatory conditions represents a key biological milestone that can be immediately translated into improvements in patient care.

**One Sentence Summary:** CD123^+^ and/or PD-L1^+^ immature and dysfunctional neutrophil subsets identified by mass cytometry, define an early human blood signature of sepsis

## Introduction

Sepsis is a life-threatening syndrome with organ dysfunction due to a dysregulated host response secondary to an infection (1). It is the 10^th^ cause of death in developed countries and the leading cause of death in the intensive care unit (ICU) (2, 3). Diagnosis of patients relies on clinical data and prognosis can be evaluated by several scores including Simplified Acute Physiology Score II (SAPS II) and Sequential Organ Failure Assessment (SOFA) Score. However, there is no robust cellular biomarker that can diagnose sepsis, distinguish it from sterile inflammation and predict its clinical outcome. A cytometry test for sepsis could provide clinically useful information in a few hours, whereas microbiological culture results typically require 18 to 72 hours and delays in treatment profoundly impact patient outcomes (4). In fact, it is estimated that the survival rate decreases by roughly 10% every hour that appropriate antimicrobial medication is delayed, emphasizing the urgent need for early diagnosis techniques (5). Other challenges in sepsis diagnosis include syndrome complexity and frequent misdiagnosis. The symptoms of sepsis overlap with both aseptic systemic inflammatory response to surgery and other noninfectious causes. Thus, a specific diagnostic test would impact patient treatment plans and could significantly improve sepsis patient outcomes. Key factors driving sepsis include the activation states of different immune cell subsets, and these cells subsets may simultaneously signal for inflammation and immune suppression in one patient (6). These cellular states are known to vary across time and patients, and thus a systems immunology approach using mass cytometry is well-suited to characterizing sepsis and related conditions in order to identify disease-specific cellular states.

Neutrophils are a primary immune cellular barrier against pathogens, but they may be a double-edged sword in sepsis. While neutrophils are usually efficient in the elimination of pathogens, their machinery can also cause local damage and organ failure (7, 8). In addition to the role of neutrophils in the inflammatory component of sepsis, neutrophils are suspected to have pathological immunosuppressive properties. Increased proportions of immature granulocytic myeloid derived suppressor cells (G-MDSC) have been associated with poor outcome (9–11). Indeed, we hypothesized that phenotype of circulating neutrophils might provide crucial early insight into immune features that drive sepsis and distinguish this disease from non-infectious inflammatory syndrome.

For the systems immunology approach here it was criticial to track features that had been identified as important in sepsis biology, but which not individually had the resolving power to specifically distinguish sepsis. Neutrophils expressing the high-affinity immunoglobulin-Fc receptor I (CD64) were described in numerous clinical studies over the last two decades (12). CD64 is normally expressed on monocytes but, in mature circulating neutrophils, its expression could be due to its upregulation during inflammation (13). High level of neutrophil CD64, measured in the peripheral blood, could also be due to released immature granulocytes from the bone marrow, especially when CD64 expression is associated with decreased expression of neutral endopeptidase (CD10) and low-affinity immuno-globulin-Fc fragment III (CD16) (9, 10, 14) (Supplemental Table1) (15). Previous studies identified also the interleukin (IL)-3 as an orchestrator of emergency myelopoiesis during sepsis, and showed its association with hospital mortality (17, 18). In parallel, programmed death ligand-1 (PD-L1) expressed on monocytes was also described as a mortality-predictor in sepsis patients (18, 19).

Accordingly, there remains an unmet need for specific and rapid diagnostic tests for sepsis, which directly reflects the immune status of sepsis patients and discriminate them from other ICU patients with aseptic inflammation. In fact, most studies that were conducted to evaluate potential cellular biomarkers specificity and sensitivity were done by comparing sepsis patients to healthy donors and by investigating one selective target at a time with limited parameters. A systems-level view is likely needed to identify cellular features that specifically distinguish sepsis infection-induced immune phenotypes from those triggered by aseptic inflammatory signals. To identify such early sepsis-specific cellular biomarkers, we developed a multi-parametric immune profiling strategy (Figure 1). Mass cytometry with a Cytometry by Time-Of-Flight (CyTOF) instrument was used to measure 42 markers on whole blood immune cells from sepsis patients and controls including healthy donnors and patients undergoing an aseptic inflammatory reaction (20). An advantage of using mass cytometry combined with computational high-dimensional data analysis is the ability to automatically and comprehensively characterize circulating immune cells, which enables identification of novel, disease-specific cellular signatures (21, 22) (Figure 1A). In fact, instead of using a classical hierarchical gating strategy we used a computational analysis approach composed of a “discovery strategy” that aims to identify sepsis-specific subsets, using a dimensionnal reduction algorithm and a clustering one (Figure 1B), and a “validation strategy”, using different set of tools (Figure 1C). Using this approach we identified two novel, early, and sepsis-specific neutrophil subsets and with an additional “expert driven strategy” we defined a small set of markers that identify these two sepsis-specific neutrophil subsets (Figure 1D), along with sepsis immune hallmarks (Figure 1E). The strenghth of this approach is the double analysis validation (computational validation and expert driven validation strategies) that confirms that the identified sepsis-specific neutrophil subsets are analysis-independent, in addition to the biological validation using a new patients’ cohort and 7-colors flow cytometry rather than multi-parametric mass cytometry that confirms the identification of these subsets are technology-independent (Figure 1F).

**Figure 1.**
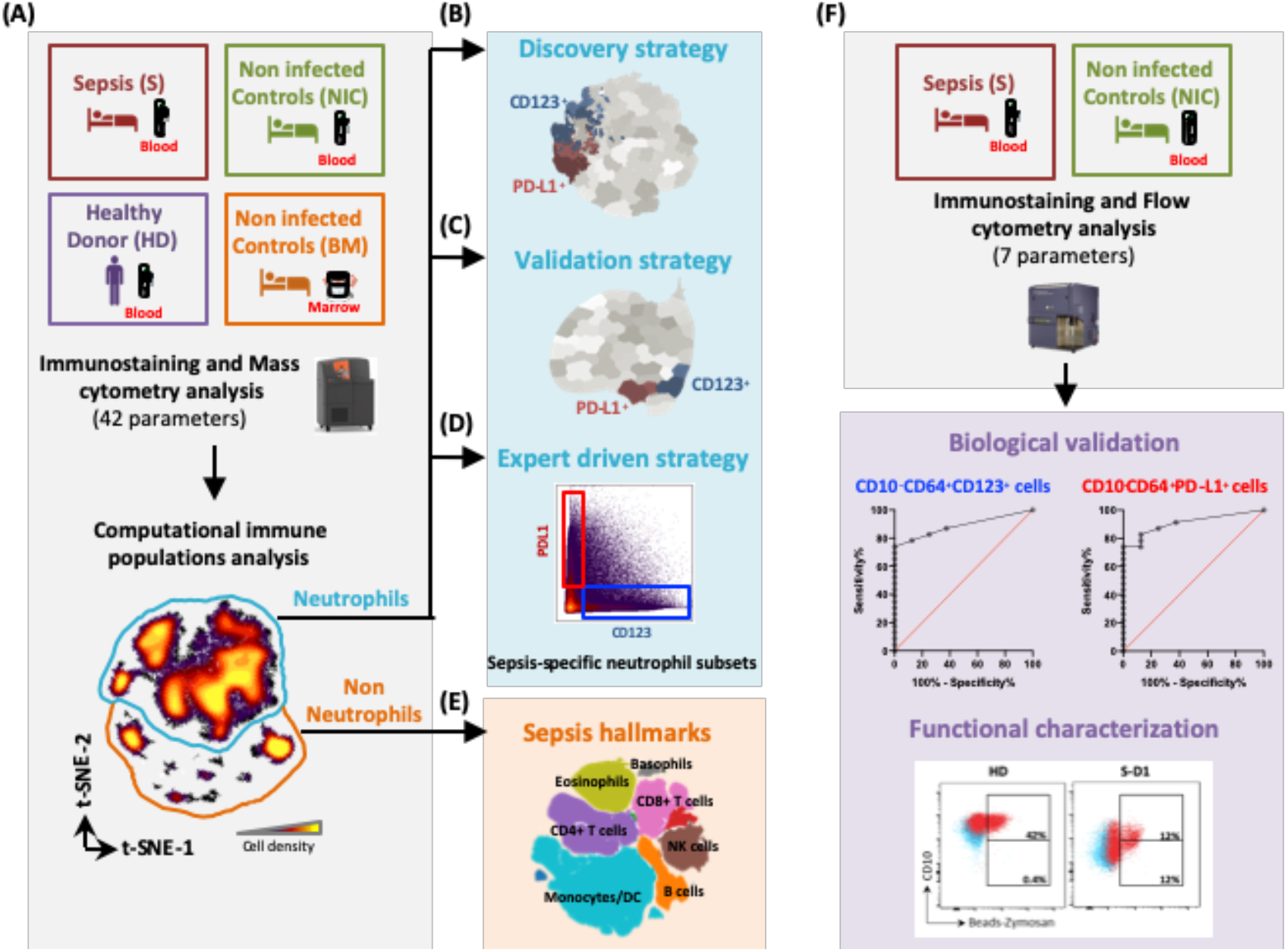
Study design. (**A**) Blood samples from sepsis patients (S) (n=17) or non-infected post-cardiothoracic surgery patients (NIC) (n=12) enrolled in the study, in addition, blood samples were obtained from healthy donors (HD) (n=11) and bone marrows biopsies from orthopedic surgery patients (BM) (n=5). Immunostainings targeting 42 parameters were performed and analyzed by mass cytometry. A computational “discovery strategy” was used to identify sepsis-specific subsets (**B**), a “validation strategy” was used to check whether the identified sepsis-specific subsets are strategy-dependant (**C**), and with an additional “expert driven strategy” we defined a small set of markers to gate on the sepsis-specific neutrophil subsets (**D**) along with the identification of sepsis immune hallmarks (**E**). A second independent cohort was used for the biological validation and functional characterization of these sepsis-specific neutrophil subsets by flow cytometry (**F**).

## Results

### Mass cytometry and computational analysis revealed sepsis-specific neutrophil signature

In an attempt to distinguish infection-induced neutrophil phenotypes from those triggered by aseptic inflammatory signals, we designed a longitudinal observational study with 40 individuals to explore the evolution of circulating immune cell phenotypes of sepsis patients (S, n=17) and non-infected post-cardiothoracic surgery patients (NIC, n=12) at day-1 and day-7 after patient’s admission in ICU (Supplemental Table2). The comparison also included blood samples from age and gender matched healthy donors (HD, n=11) and bone marrow biopsies from orthopaedic surgery patients (BM, n=5) (Figure 1A). Whole blood immunostaining was performed, within 3h after blood drawing, using a customized, 42-parameter mass cytometry panel designed to give a comprehensive evaluation of circulating leukocytes (Figure 1A, Supplemental Table 3). We then combined high-dimensional mass cytometry with computational data analysis to automatically identify circulating immune cell population. Neutrophils were automatically separated and gated appart from other circulating immune cells, using Visualization of t-Distributed Stochastic Neighbor Embedding (viSNE implementation of t-SNE) (23) (Figure 1A), to allow for independent analysis downstream (Figure 1E).

The neutrophils were analyzed using a “discovery starategy” based on a dimentionality reduction with t-SNE followed by a clustering of the cells in nodes with Spanning-tree Progression Analysis of Density-normalized Events (SPADE) (24). This strategy allowed to define an imprint for each sample group. In an unsupervised manner, S day-1 patients’ blood neutrophils were arranged in the upper left area of the map whereas the NIC patients’ cells were in the central/upper right area and HD neutrophils were located on the lower right area of the map (Figure 2A). These S neutrophils were clustered in specific nodes that were absent from NIC and HD (Supplemental Figure 1, 2) and some of these S specific nodes were shared with BM, suggesting occurrence of myelocytosis for S patients (Supplemental Figure 1, 2). Most cells from day-7 samples were phenotypically similar to samples from HD (Figure 2A, Supplemental Figure 1, 2). CD16, CD10 and CD64 markers split neutrophils signature into two positive and negative subpopulations for each marker (Supplemental Figure 1). To identify the phenotype of S-specific neutrophil nodes, sample and node abundance were arranged according to complete linkage hierarchical clustering (Figure 2B, Supplemental Figure 3), and the phenotype of each node was presented in Figure 2C (Supplemental Figure 4). Most of the samples of each patients’ group were clustered together with the exception of three S day-1 patients (pink) and two NIC day-1 samples (green). S day-1 and BM (orange) samples were clustered together. Day-7 samples were more scattered, with half of them close to HD suggesting an immune recovery. This strategy allowed the precise delimitation of four groups of cell nodes (Figure 2D): ^(1)^ HD-abundant nodes representing neutrophils with high expression of CD16, high to mid expression of CD10 and no expression of CD64, ^(2)^ NIC and S day-7 common nodes harbouring a CD16 low to high expression, CD10 mid expression and no expression of CD64, ^(3)^ day-1 NIC and S common nodes defined as CD16^low^CD10^−^CD64^low^, and ^(4)^ S day-1 and BM nodes with a high to low expression of CD64 and no expression of CD10. The main phenotypic differences observed in groups of nodes ^(4)^ were the level of expression of CD123 or PD-L1 (Figure 2C, D). On this basis, meta-clusters were generated to group nodes that share similar phenotype (Supplemental Figure 5A) and that statistically discriminate S and NIC at day-1 (Supplemental Figure 5B). Two novel subsets specific to S at day-1 were identified and observed to be relatively lacking in NIC neutrophils (Figure 2E). The first subset (in red) was composed of CD10^−^CD64^+^CD16^+^PD-L1^+^ neutrophils (nodes 11, 38, 50 and 54) (S median proportion: 18.08±23.33 %, NIC median proportion: 0.81±2.92 %, *p*=*0.0002*) and the second one (in blue) identified as CD10^−^CD64^+^CD16^low^CD123^+^ immature neutrophils (nodes 10, 27 and 35) (S median proportion: 10.06±23.76 %, NIC median proportion: 0.04±0.90 %, *p*<*0.0001*) (Figure 2E). We also recapitulated previously described results regarding the sepsis related increase of circulating immature CD10^−^CD64^+^ neutrophils (nodes 16, 24, 41, 49, and 55) when compared to NIC at day-1 (S median proportion: 11.03±21.05 %, NIC median proportion: 0.62±4.69 %, *p*=*0.001*) and we confirmed their phenotypic similarities with a third of BM neutrophils (BM median proportion: 37.39±14.75 %) (9, 10, 14) (Figure 2E). Also, we noticed that all HD specific-nodes were absent in S patient day-1 samples (Figure 2B, D).

**Figure 2.**
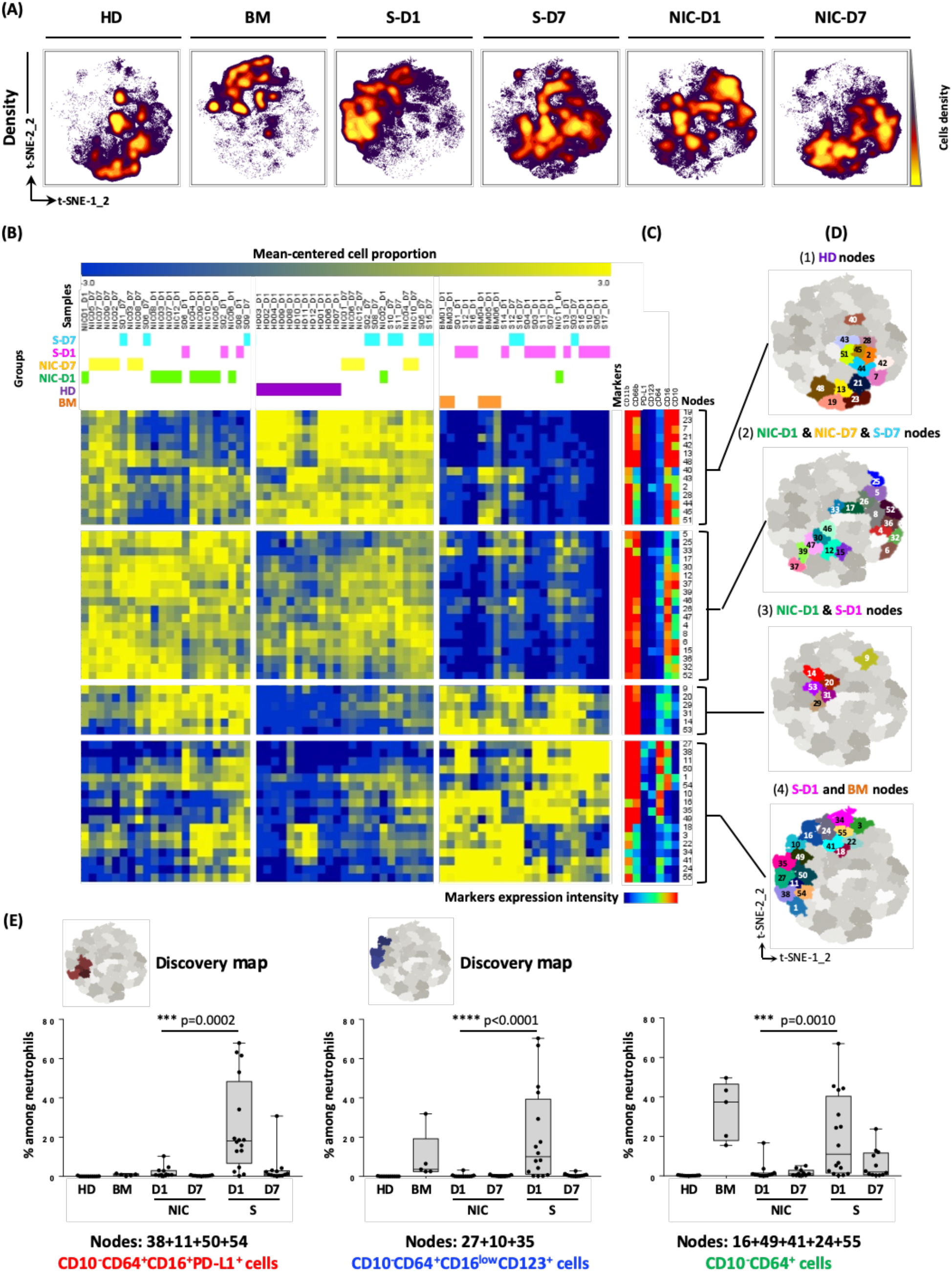
Identification of sepsis day 1-specific neutrophils with a discovery analysis strategy. **(A)** t-SNE analysis was performed on neutrophils from all samples with cells being organized along t-SNE-1-2 and t-SNE-2-2 according to per-cell expression of CD11b, CD66b, CD16, CD10, CD64 and CD123, PD-L1. Cell density for the concatenated file of each group is shown, on a black to yellow heat scale, for each group time-point. (**B**) A heat map shows samples clustering (columns) according to nodes cell proportion log2-transformed and centered around the mean proportion of all samples’ nodes (rows). Samples and mean-centered log2-transformed nodes cell proportion were arranged according to complete linkage hierarchical clustering. Heat intensity (from blue to yellow) reflects the mean-centered log2-transformed cell proportion of each sample’s node. (**C**) A heatmap shows characterization of cell nodes identified by SPADE (columns) according to mean expression of 7 markers (rows). Markers were arranged according to complete linkage hierarchical clustering and nodes were pre-ordered according to (**B**) heat map nodes order. Heat intensity (from blue to red) reflects the mean expression of each marker for each node. (**D**) Four groups of nodes were back-viewed on t-SNE1-2 / t-SNE2-2 map. (**E**) cells abundance of each meta-cluster subset (CD10-CD64+CD16+PD-L1+ cell subset in red, CD10-CD64+CD16lowCD123+ cell subset in blue and CD10-CD64+ cell subset in green) was presented as cell proportion among total neutrophils of each group samples. Statistics: Nonparametric two-tailed Mann-Whitney test was used to compare differences in cellular abundance of cell subsets between NIC-D1 and S-D1 (see the Methods section). Sample sizes: HD=12, BM=5, NIC-D1=12, NIC-D7=11, S-D1=16 and S-D7=12.

With this strategy, two novel neutrophil subsets were identified, including CD123^+^ cells (red) and PD-L1^+^ cells (blue), and the absence of HD neutrophil phenotypes at an early stage of sepsis.

### A computational validation strategy confirmed sepsis day-1 specific neutrophil subsets

To test whether the previously identified neutrophil subsets were sepsis-specific and strategy-independent, a second unsupervised data analysis strategy was applied on the same files used in the discovery strategy (Figure 1C).

The “validation strategy” based on a dimentionality reduction with Uniform Manifold Approximation and Projection (UMAP) (25) followed by a clustering of the cells with the FlowSOM algorithm (26) showed cells from the different groups in different areas of the low-dimensional space (Supplemental Figure 6A). This strategy allowed the identification of 50 neutrophil clusters and the complete linkage hierarchical clustering of their relative cell abundance arranged again with the samples of each patients’ group together (Supplemental Figure 6B); with S day-1 patients (pink) and NIC day-1 samples (green) at the extremities of the dendrogram. Two main cell cluster groups (pink gates) appeared to be more abundant in sepsis samples (Supplemental Figure 6B, C) and almost all HD associated-clusters (purple gate) were absent in sepsis patient day-1 samples.

To phenotypically characterize the pink gate clusters, Marker Enrichment Modeling (MEM) (27, 28) was used. The MEM label of each cluster is an objective description of what makes that subset distinct from the other 50 clusters. Among these clusters, three groups of clusters were identified, the group of CD10^−^CD64^+^ immature cells (pink clusters), and two clusters groups phenotypically identical to the “discovery strategy” sepsis-specific neutrophils nodes (Supplemental Figure 6D, Figure 3A). Clusters 18 and 19 (in red) contained CD10^−^CD64^+^PD-L1^+^ neutrophils with a median cell proportion of 5.50±29.83 % for S day-1 samples and 0.09±1.15 % median proportion for NIC day-1 samples (*p*<*0.0001*) (Figure 3B). Clusters 6 and 7 (in blue) gathered CD10^−^CD64^+^CD16^low/−^ CD123^+^ immature neutrophils with median cell proportions of 2.43±17.81 % and 0.04±1.14 % for S day-1 and NIC day-1 samples respectively (*p*=*0.0006*) (Figure 3B). We also visually noted that red clusters (PD-L1^+^ cells) and blue clusters (CD123^+^ cells) from the “validation strategy” co-localised with red nodes (PD-L1^+^ cells) and blue nodes (CD123^+^ cells), respectively, from the “discovery strategy”, when back mapped onto the t-SNE1-2 / t-SNE2-2 axes (Figure 3C).

**Figure 3.**
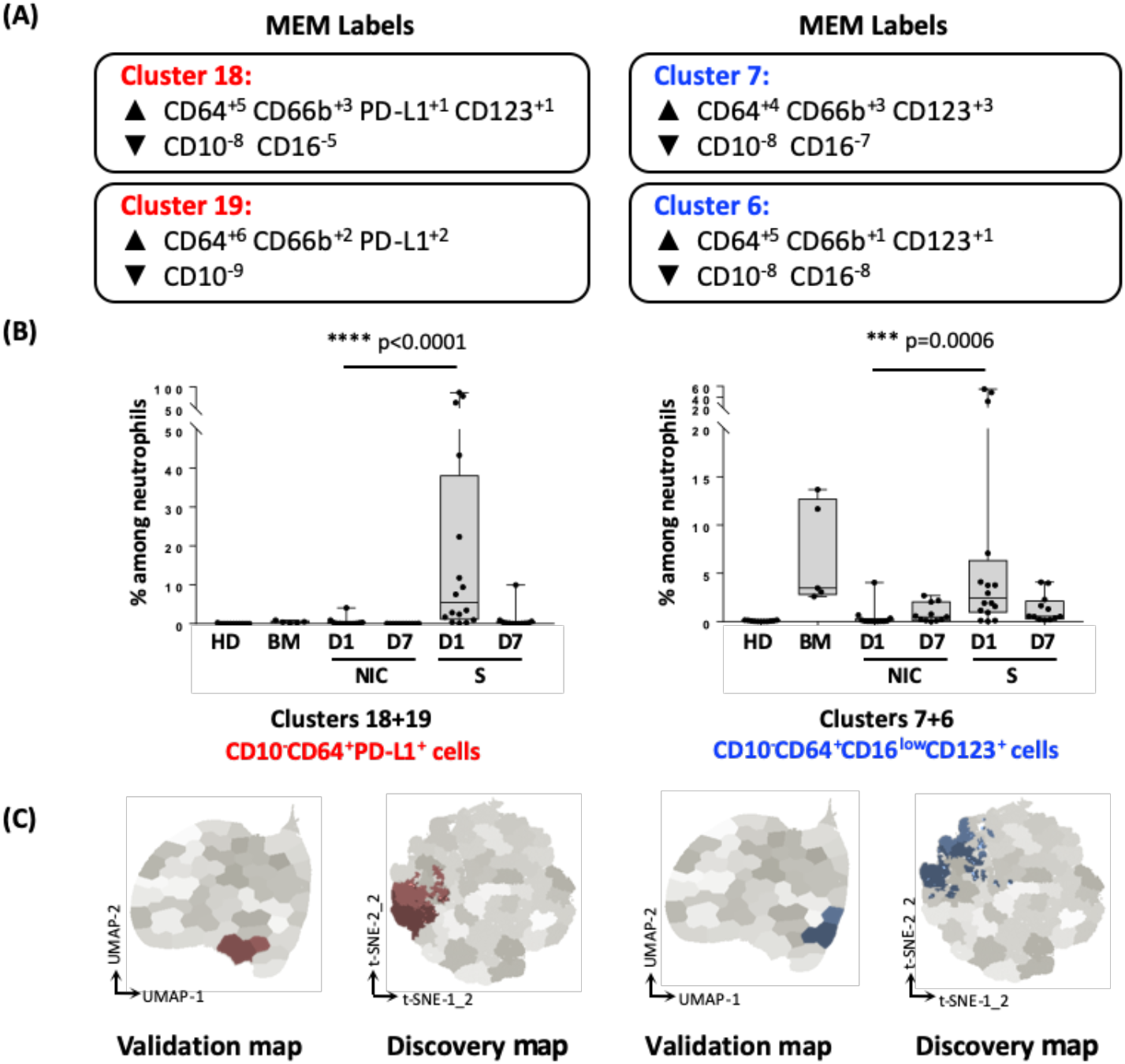
Validation of sepsis day-1-specific neutrophil subsets by a second computational strategy. As a first step, UMAP analysis was performed on all samples neutrophils and cells were organized along UMAP-1 and UMAP-2 axes according to per-cell expression of CD11b, CD66b, CD16, CD10, CD64 and CD123, PD-L1. As a second step, FlowSOM clustering was done to separate neutrophils subsets into 50 clusters. MEM was then used to quantify the enriched features of the 50 clusters. Protein enrichment was reported on a +10 to −10 scale, where +10 indicates that protein’s expression was especially enriched and −10 indicated that the protein’s expression was excluded from those cells, relative to the other neutrophils clusters. (**A**) Among these clusters, two meta-clusters were identified as phenotypically identical to the strategy-1 sepsis-specific neutrophils: clusters 18 and 19 (in red) composed of CD10-CD64+PD-L1+ neutrophils and clusters 6 and 7 (in blue) composed of CD10-CD64+ CD16lowCD123+ neutrophils. (**B**) Cells abundance of each meta-cluster subset (CD10-CD64+CD16+PD-L1+ cell subset in red and CD10-CD64+CD16lowCD123+ cell subset in blue) was presented as cell proportion among total neutrophils of each group samples. Statistics: Nonparametric two-tailed Mann-Whitney test was used to compare differences in cellular abundance of cell subsets between NIC-D1 and S-D1 (see the Methods section). Sample sizes: HD=12, BM=5, NIC-D1=12, NIC-D7=11, S-D1=16 and S-D7=12. (**C**) each meta-cluster cells (red and blue) was back-viewed on both UMAP-1 / UMAP-2 map, and t-SNE1-2 / t-SNE2-2 map.

### Expert gating strategy with a small set of markers validated sepsis day-1 neutrophil signature that correlate with SAPSII and SOFA scores

After cell subsets were identified by automatic and high-dimensional analysis strategies, we determined whether the identified neutrophil signature could be found using conventional analysis applicable by experts. The use of such gating strategy would make it easier to transpose it to clinical use.

A bi-parametric gating strategy on a limited set of markers allowed the identification of neutrophils expressing CD123 and PD-L1 (Figure 4A). When CD123^+^ and PD-L1^+^ sepsis-specific neutrophils were mapped back onto both t-SNE1-2 / t-SNE2-2 axes and UMAP1 / UMAP2 axes, they located in the same regions as the cells identified by the two previous computational strategies meaning that they share the same phenotype (Figure 4A). This expert gating strategy applied on the current dataset, allowed the selection of PD-L1^+^ neutrophils that were significantly more abundant in S patients’ blood collected at D1 post-admission to ICU (9.25±27.02 %) when compared to day 1 non-infected post-cardiothoracic surgery patients (NIC) (0.12±0.37 %, p<0.0001) or HD (0.01±0.03 %, p<0.001) (Figure 4B). Similarly, expert gating allowed the selection of S-specific neutrophils (2.47±17.55 %) that were consistent with CD123^+^ red subsets cells phenotype and that were almost absent from day 1 NIC (0.04±0.47 %, p<0.0001) or HD (0.04±0.04 %, p<0.0001) (Figure 4B). Unlike PD-L1^+^ sepsis-specific neutrophils, the proportion of CD123^+^ sepsis-specific, assessed by the simple gating strategy on mass cytometry data, correlate positively with SAPS II score (r=0.62, *p*=0.0077, R^2^=0.46) and with SOFA score (r=0.55, *p*=0.0037, R^2^=0.31) (Figure 4C).

**Figure 4.**
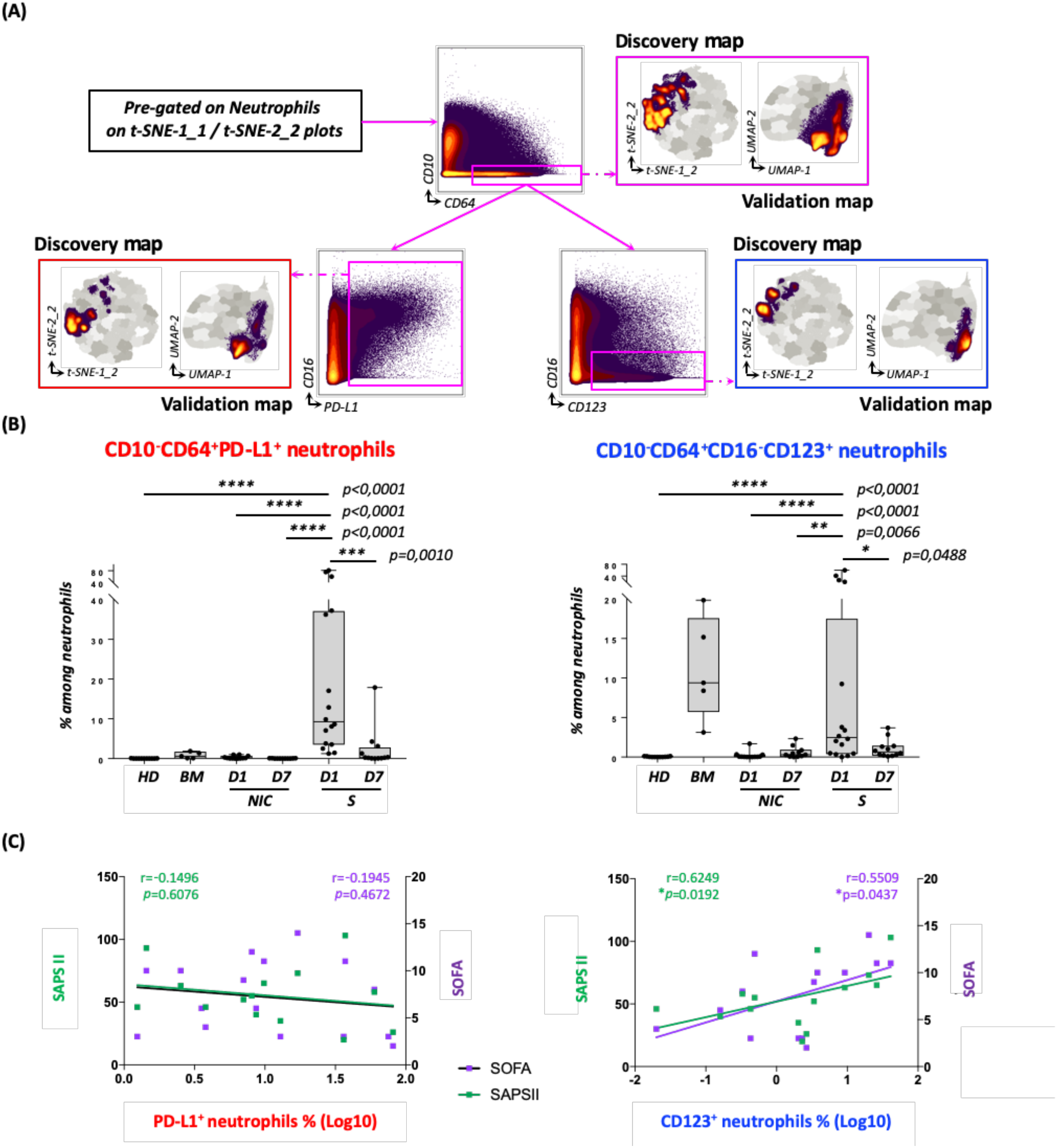
Sepsis day 1-specific neutrophil subsets validated by expert gating correlate with severity scores. Expert gating strategy with 7 markers set (**A**) allowed the selection of CD10-CD64+PD-L1+ cell subset (in red) and CD10-CD64+CD16lowCD123+ cell subset (in blue), back-viewed on both UMAP-1 / UMAP-2 map, and t-SNE1-2 / t-SNE2-2 map. The two neutrophil subsets are significantly more abundant in sepsis patients (S) blood collected at day-1 post-admission to ICU when compared to day-1 or day-7 non-infected post-cardiothoracic surgery patients (NIC) or Healthy donors (HD) (**B**). Correlation between the log10 transformed frequency of CD10^−^CD64^+^PD-L1^+^ neutrophils subset (in red) or CD10^−^ CD64^+^CD16^low^CD123^+^ neutrophils subset (in blue) and SAPS II score (green dots) or SOFA score (purple square) are shown in (**C**). Statistics: Nonparametric two-tailed Mann-Whitney test was used to compare differences in cellular abundance of cell subsets between S-D1 and NIC-D1, NIC-D7 or HD and nonparametric two-tailed Wilcoxon signed-rank test was used to compare differences between the two matched groups S-D1 and S-D7. Linear regression curves, and Spearman correlation test spearman were used to assess correlation between neutrophil subsets frequency and severity scores (see the Methods section). Sample sizes: HD=12, BM=5, NIC-D1=12, NIC-D7=11, S-D1=16 and S-D7=12.

Thus, a simple gating strategy assessing only 7 key markers identified successfully CD123^+^ and PD-L1^+^ sepsis-specific neutrophils and indicated that CD123^+^ neutrophils may be a marker of sepsis severity and could be considered a prognostic indicator.

### Mass cytometry and unsupervised analysis identified classical sepsis immune hallmarks

In order to identify an early and globale sepsis-specific immune signature, that may reinforce the CD123^+^ and PD-L1^+^ neutrophil subsets as sepsis biomarker candidates, the non-neutrophils circulating immune cells were computationaly analyzed using t-SNE and SPADE algorithms. A heatmap was generated to characterize nodes phenotype and to delimitate the main circulating non-neutrophil immune cell populations, according to complete linkage hierarchical clustering (Supplemental Figure 7A). These populations were then color coded and backgated on the t-SNE map (Supplemental Figure 7B). Classical hallmarks of sepsis were identified, including lymphopenia, monocytopenia and a persistent lower level of monocytes HLA-DR in S patients when compared to HD group (*p*<0.0001, *p*=0.0426 and *p*<0.0001 respectively, Figure 5A). In parallel to these results, we also observed an elevated number of circulating neutrophils *(p*=0.0039), and consistent with that, a higher neutrophil to lymphocyte ratio (*p*<0.0001) in S vs. HD (Figure 5B). These trends were not exclusive to S, but were also observed in NIC group when compared to HD group (*p*=0.0003, *p*<0.0001, *p*=0.0034, *p*<0.0001, for lymphocytes and neutrophils counts, monocytes HLA-DR expression level and neutrophils/lymphocytes ratio, respectively). No significant difference was observed between S and NIC group at day-1 within these main immune cell populations (Supplemental Table 2).

**Figure 5.**
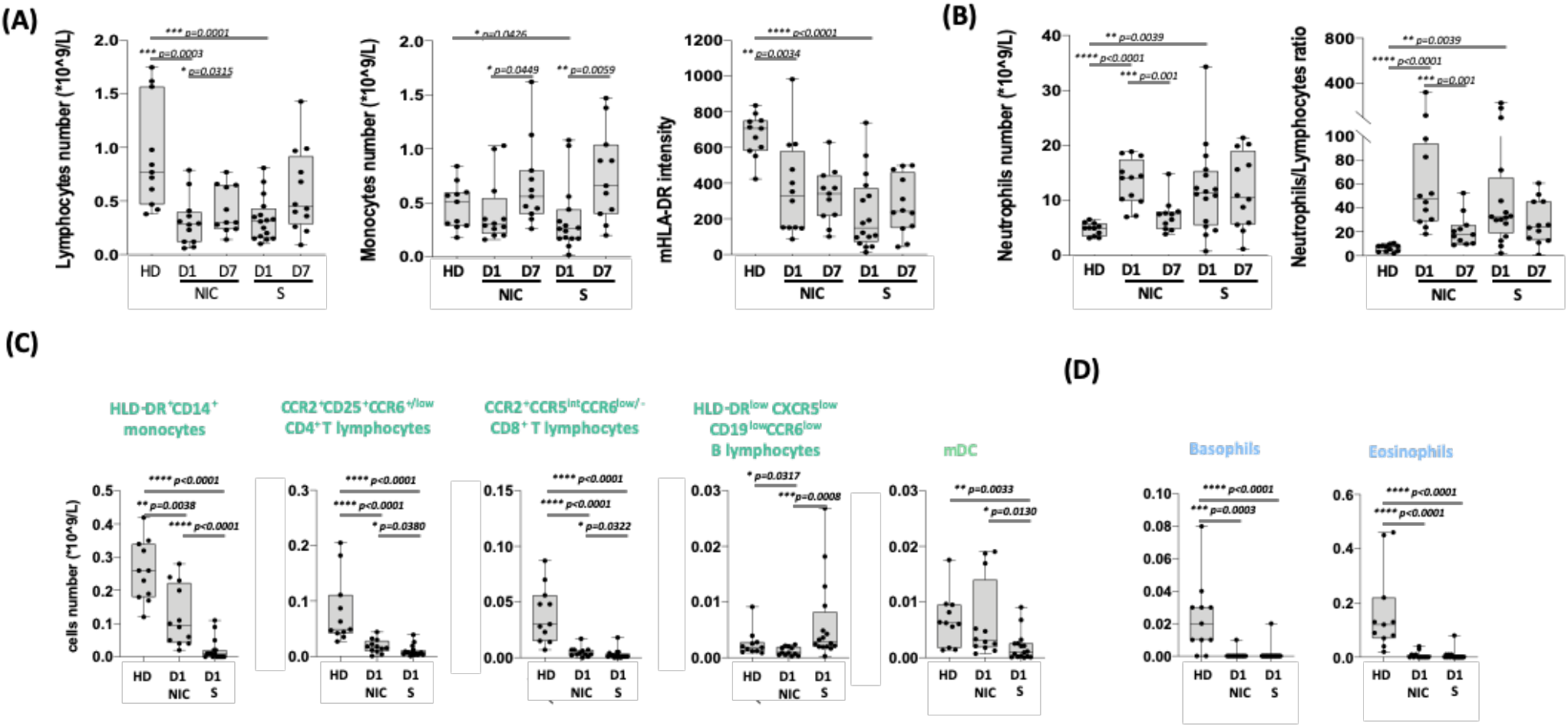
Non-neutrophil cells analysis resume sepsis immune hallmarks. **(A)** Lymphocytes and monocytes numbers and intensity of HLA-DR expression on monocytes (mHLA-DR) were obtained from non-neutrophils computational analysis and presented for each group. **(D)** Neutrophils numbers were obtained previously from the computational separation of neutrophils from non-neutrophil cells and used to calculate Neutrophils/Lymphocytes ratio. Cell number of the main immune subsets that were differentially abundant in S group from HD and NIC were presented in **(C)** and the ones that were differentially abundant in S group from only HD were presented in **(D)**.

To identify an early sepsis-specific signature within these immune populations, we compared the abundance of the identified cell nodes of these immune populations between HD, NIC and S samples at day-1. The abundance of 22 nodes was found selectively regulated in S at day-1 when compared to both NIC and HD and 25 nodes differentiated S only from NIC at day-1 (Supplemental Figure 7C, D). It included notably 15 nodes identifying classical monocytes with high expression of HLD-DR, 3 nodes of CD4^+^ T lymphocytes and CD8^+^ T lymphocytes expressing CCR2 and CCR6, all were highly reduced in S patients, one node of B lymphocytes with a low expression of B cells pan markers (HLD-DR, CXCR5, CD19 and CCR6) and one node identified monocyte-derived DC (Figure 5C). Among the nodes that were massively reduced in both S and NIC sample, 15 nodes out of 55 represent Basophils and Eosinophils subsets (Figure 5D); the others being scattered among other cell populations.

Taken globally, the analysis of circulating non-neutrophil cells with a computational strategy allowed us to resume sepsis hallmarks and identify the differences of several circulating immune subsets abundance with no added value to the identified sepsis-specific neutrophils.

### CD123+ and PD-L1+ sepsis-specific neutrophils are detectable by conventional cytometry and discriminate infected and non-infected patients

To evaluate the efficiency and specificity of CD123^+^ and PD-L1^+^ neutrophil subsets to discriminate sepsis patients from non-infected ones, we set up a fluorescent flow cytometry panel composed of 7 surface markers (Supplemental Table 4) that we used to monitor a new validation cohort composed of non-infected patients (n=8) and sepsis patients (n=23) independent from the previously described ones.

With the overlay of full minus-two (FMT) stained control and the full panel stained tubes of three representative patients of several expression levels of CD10, CD123 and PDL1, we can appreciate the increase of CD123^+^ and PD-L1^+^ sepsis-specific neutrophil subsets with the decrease of CD10 expression by neutrophils (CD14^−^CRTH2^−^CD15^+^ cells) (Figure 6A).

**Figure 6.**
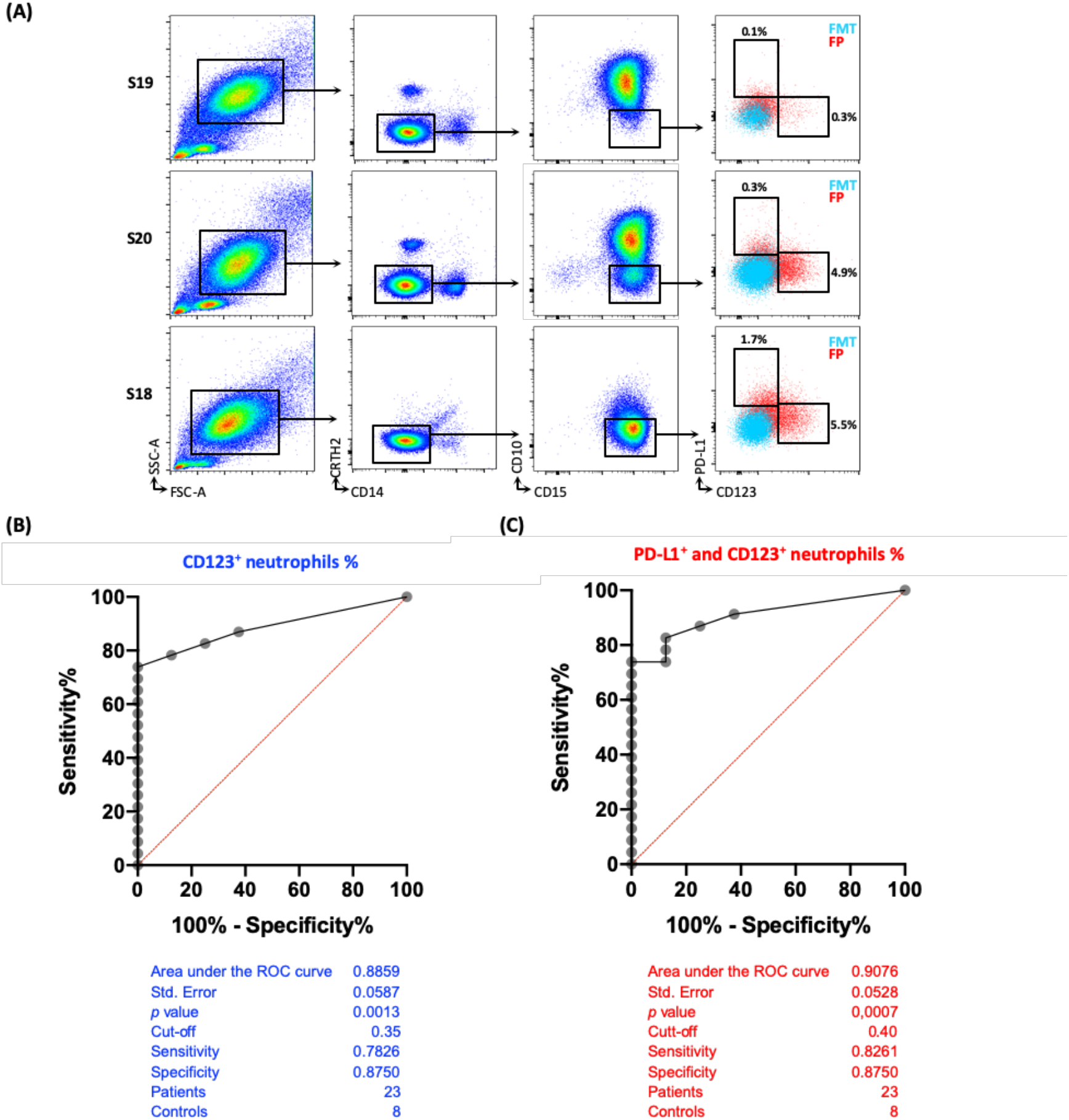
Sepsis-specific neutrophils are detectable by conventional cytometry and discriminate infected from non-infected patients. The gating strategy applied on fluorescent flow cytometry data of three new sepsis patients is showed in (**A**). The overlay of full minus-two (FMT) stained control and the full panel (FP) stained tubes of each representative patient, showed the increase of sepsis-specific neutrophil subsets with the decrease of CD10 expression by neutrophils (CD14-CRTH2-CD15+ cells). The ROC curves obtained using CD123^+^ neutrophil subset or CD123^+^ and PD-L1^+^ neutrophil subsets were presented in (**B**) and **(C)** respectively.

Based on flow cytometry results, a CD123^+^ neutrophil subset proportion cut-off point of 0.35 %, was able to rule out sepsis patients with a specificity of 87.50% and sensitivity of 78.26% (AUC of 0.89 with *p*=0.0013) (Figure 6B). When combining both CD123 and PD-L1 neutrophil subsets proportions and a cut-off point of 0.40%, the increase in the sensitivity was modest (sensitivity of 0.83 (*p*=0.0007) without any effect on the specificity (Figure 6C).

These data indicated that these subsets could be reliably quantified by traditional clinical flow cytometric profiling and may help in patients diagnosis.

### Immature sepsis neutrophils exhibit an impaired microbial specific activation and phagocytosis

To address sepsis immature (CD64+CD10^−^) neutrophils activation and phagocytic capacities, whole blood of each tested individual was incubated with *Staphylococcus aureus* (*S. aureus*) or Zymosan coated bio-particles labelled with pHrodo, a pH-sensitive fluorochrome in order to evaluate phagocytic competent cells (29). After 1h incubation at +37°C for the positive controls (PC) or in ice for the negative controls (NC), cells were analyzed by flow cytometry to identify immature neutrophils bio-particles uptake and cells activation. All circulating neutrophils were able to phagocyte *S. aureus* beads independently from their group (HD, S-D1, BM). However, S day-1 neutrophils phagocytosis of Zymosan Beads (Mean±SD = 28.12±8.39%) was not as effective as that of HD (Mean±SD = 50.43±13.04, *p*=0.02) (Figure 7A). t-SNE visualisation of PC and NC neutrophils of both *S. aureus* (Figure 7B) and Zymosan (Figure 7C) bead stimulations highlighted the lower expression level of CD11b marker by S day-1 neutrophils when compared to HD and the default of activation of these cells after microbial beads activation. In fact, S neutrophils exhibited a lower ratio of CD11b and CD66b MFI between PC and NC after activation, when compared to healthy donors after *S. aureus* (Figure 7D) or Zymosan (Figure 7E) stimulations. The impaired phagocytic capacity of sepsis-patients’ immature neutrophils compared to HD neutrophils was confirmed by the measurement of phagocytosed beads MFI ratios between PC and NC. This ratio was three times lower for S day-1 *S. aureus* response (Figure 7D) and 30% lower for S day-1 Zymosan response (Figure 7E). This part of the study allowed the identification of an impaired capacity of immature sepsis neutrophils to form efficient phagolysosomes after bio-particles stimulation and a default of activation when compared to HD.

**Figure 7.**
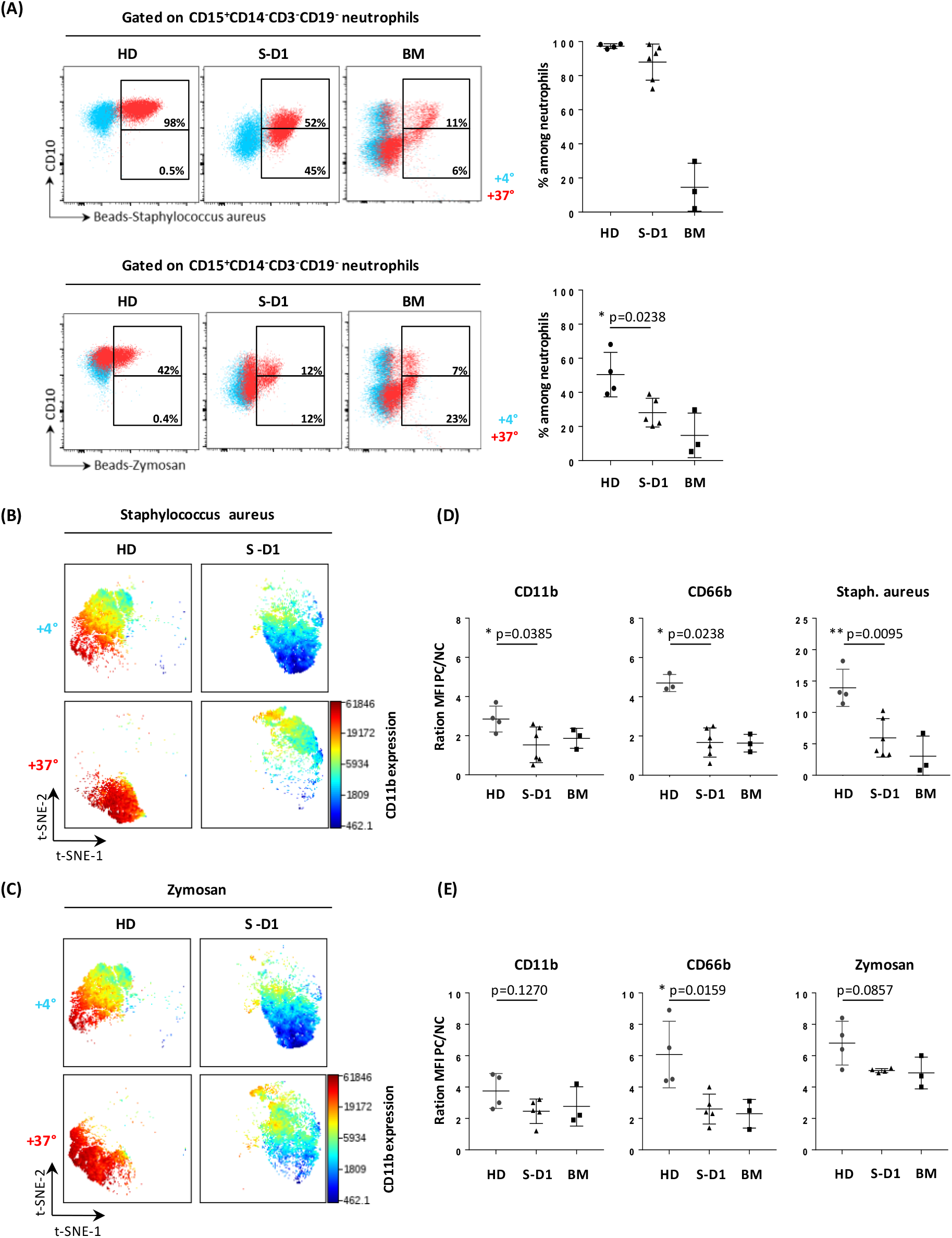
Staphylococcus aureus and Zymosan specific activation and phagocytosis are impaired in immature sepsis neutrophils. To address sepsis immature (CD64+CD10−) neutrophils phagocytic capacities, 100μL of blood were incubated with 20μL or 40μL of beads coated with Staphylococcus aureus or Zymosan, respectively, coated-particles and coupled with pH acidification-sensitive fluorochrome. After 1h incubation at 37°C (PC: positive control) or 4°C (NC: negative control) cells were stained and analyzed by flow cytometry. (**A**) represents gating strategy of CD15+CD14−CD3−CD19− neutrophils from healthy donors (HD), sepsis day-1 samples (S-D1) and bone marrow samples (BM). Cells were separated in 2 gates based on CD10 expression and phagocytosis marker intensity (Staphylococcus aureus or Zymosan) and cells from PC (red dots) were overlaid on NC cells (blue dots). The proportion of total phagocytic neutrophils were presented for the three groups. t-SNE analysis organized cells along t-SNE axes according to per-cell expression of 5 proteins and phagocytosis fluorescence. Cell expression of CD11b after Staphylococcus aureus (**B**) or Zymosan (**C**) stimulations, for one representative individual of HD and S-D1 stimulated at +4°C (NC) and +37°C (PC) is shown on a heat scale. The ratio between PC and NC CD66b CD11b and particles MFI, of each individual after Staphylococcus aureus (**D**) or Zymosan (**E**) stimulations, in each group were plotted in histograms. CD10-cells have less phagocytic capacity whatever it is appreciated by MFI or proportion. Stimulated CD10-cells exhibit a lower level of expression of CD11b and CD66b. Statistics: Nonparametric two-tailed Mann-Whitney test was used to compare differences in cellular abundance of cell subsets and MFI ratios (see the Methods section). Sample sizes: HD=4, S-D1=6 and BM=3.

## Discussion

Stratification of patients on the basis of cellular biomarkers is an unmet need in sepsis patient care. Here, the goal was to identify cellular signatures that reflect directly the immune status of sepsis patients and to discriminate sepsis patients from other ICU patients with aseptic inflammation. Whole blood mass cytometry and unsupervised analysis identified classical hallmarks of sepsis, including lymphopenia and an elevated amount of circulating neutrophils, and revealed two novel neutrophil subsets that distinguish early sepsis from aseptic inflammatory syndromes. Previous studies in sepsis evaluated only few markers simultaneously and only on enriched cell populations from peripheral blood mononuclear cells (PBMC). CyTOF high-dimensional technology coupled with two unsupervised analytical strategies were used to pinpoint what immunologically discriminates, in whole blood, infection from inflammation and the result of these automated strategies was confirmed by a traditional expert gating strategy and by classical flow cytometry. Two novel neutrophil subsets were identified, CD10^−^CD64^+^PD-L1^+^ neutrophils and CD10^−^ CD64^+^CD16^low/−^CD123^+^ immature neutrophils that could be used for sepsis diagnosis. Furthermore, CD123 subset correlates with prognosis scores.

The results of this study recapitulated previous findings regarding the sepsis-related increase of circulating immature CD10^−^CD64^+^ neutrophils (9, 10, 14). Mass cytometry identified classical hallmarks of sepsis, including lymphopenia, monocyte HLA-DR downmodulation, and a higher amount of neutrophils with a phenotype that was distinct from HD. In this work, the focus was on neutrophils that were previously targeted in several studies as key biomarker cells for sepsis. In a couple of studies, the authors proposed staining kits or automated analysis strategies to evaluate CD64 level in patients’ circulating neutrophils by flow cytometry (30, 31) and in others, microfluidic systems were proposed as point-of-care methods for CD64 quantification (31, 32). Several meta-analysis studies have also shown that, particularly when SOFA score is combined with CD64^+^ neutrophils, the accuracy, sensitivity and specificity of sepsis diagnosis at early stages of disease can be dramatically improved (33). Despite all these large efforts, the CD64 detection-based tools are not yet standardized for sepsis diagnosis, because of the heterogeneity of sepsis syndrome and inter-individual variability of CD64 basal level among sepsis patients.

The CD10^−^CD64^+^CD16^low/−^CD123^+^ population is most consistent with immature neutrophils. The frequency of this population among total neutrophils positively correlates with both SAPS II and SOFA severity scores used in clinical practice for sepsis prognosis, and need to be confirmed in a larger collection. The expression of CD123 receptor by neutrophils was not described before during sepsis. In a previous study of Weber *et al.,* using a mouse model of abdominal sepsis, the cytokine IL-3 was reported to potentiate inflammation in sepsis by inducing myelopoiesis of neutrophils and IL-3 deficiency protects mice against sepsis (16). Moreover, the authors described an association between high plasma IL-3 levels and high mortality. This result was also obtained in a recent prospective cohort study, where higher levels of IL-3 were shown to be independently associated with hospital mortality in septic patients (17). All these results identify IL-3 and its receptor CD123 as an orchestrator of emergency myelopoiesis, and reveals a new target for the diagnosis and treatment of sepsis.

To our knowledge, the expression of PD-L1 by neutrophil during sepsis was not reported before. It was defined on monocytes, macrophages and endothelial cells (34) but not granulocytes. Monocyte PD-L1 expression was described as an independent predictor of 28-day mortality in patients with septic shock (18, 19). Peripheral blood transcriptomic analysis done by Uhle *et al*., revealed the expression of PD-L1-gene among the top 44 immune-related genes differentially expressed between patients with sepsis and healthy donors (11). In parallel, mice in which the PD-1/PD-L1 interaction was inhibited show improved survival to sepsis (35). Our results bring up a new target for the immune checkpoint therapies that block T cell exhaustion PD-1/PD-L1 axis and that were proposed as new candidate to treat sepsis patients.

Controversial results were previously described regarding functional aspects of neutrophils during sepsis. On one hand, Demaret *et al*., described conserved phagocytosis and activation capacities of sepsis neutrophils characterized as CD10^dim^CD16^dim^ immature cells, after whole blood IL8, fMLP or FITC-labeled *Escherichia coli* stimulation cells (36). On the other hand, Drifte *et al*., by comparing mature and immature neutrophils functions found that the latter were less efficient in phagocytosis and killing. Accordingly, we observed an impaired capacity of cells to form efficient phagolysosomes after bio-particles stimulation and a default of activation when compared to HD.

The immunosuppressive function was also attributed to G-MDSC neutrophils subset during sepsis (9–11, 14). But, to date, human G-MDSC definition lacks consensual phenotypic characterization. Published results on G-MDSC in cancer were obtained according to various phenotypes. Condamine *et al.* described them as Lectin-type oxidized LDL receptor-1 (LOX1) expressing cells (37). Using flow cytometry, we measured the expression of LOX-1 in sepsis patients (data not shown). No LOX-1 co-staining was observed with neither CD123^+^ nor PD-L1^+^ subsets. More investigation is needed to characterize if CD123^+^ neutrophils and PD-L1^+^ subset belong to G-MDSC.

Further research should be conducted to identify appropriate clinical actions for each identified neutrophil subset, to understand whether altered neutrophil production is responsible for increased sepsis risk, and to determine how these subsets can be therapeutically targeted.

In parallel, future studies should now be undertaken to validate the use of these new neutrophil subsets in clinic by routine flow cytometry as an early biomarker predictive of sepsis. Larger cohorts that better represent not only sepsis patients but also the diversity of aseptic inflammatory syndromes need to be evaluated. The use of a whole blood flow cytometry test to diagnose sepsis could change the fate of patient’s care. The clinician would have a rapid and specific result, obtained before microbiological cultures results, that could guide their therapeutic decision.

## Methods

### Study design

This observational study was approved by the Comité de Protection des Personne Paris VII ethic committee (CPP IDF VII A000142-53) which waived the need for written informed consent since it was with low risk for the patients and with no need for specific procedure besides routine blood sampling. The study was conducted in 4 ICUs: 2 surgical ICUs (Hôpital Lariboisière, Hôpital Saint Louis), 1 medical ICU (Hôpital Lariboisière), and 1 post-cardiac surgery ICU (Hôpital Europeen Georges Pompidou). All individuals included in the study were adults (>18 years of age) with no pre-existent immunosuppression history.

Seventeen sepsis (S) patients (10 male and 7 female), with a median of age of 75 years old were included in the study (Supplemental Table 2). Sepsis was defined in accord to The Third International Consensus Definitions for Sepsis (4), i.e. an increase of 2 or more points on SOFA score in related to a suspected or confirmed infection. All patients received prompt treatment with antibiotics, adequate fluid resuscitation, and support for damaged organ function, including mechanical ventilation and continuous renal replacement therapy. Twelve patients (8 male and 4 female), with a median of age of 75, undergoing non-infection related cardiac intervention requiring cardiopulmonary bypass were also included in the study. They were considered in this study as non-infected inflammatory controls (NIC).

Patients’ clinical data were collected at day 1 and day 7 and included vital status, component of SAPS II and SOFA score, clinical and biological parameters and blood count. Vital status was checked at day 28 (Supplemental Table 2). Blood samples drawn in heparin-coated tubes, were collected one and seven days post admission of sepsis patients or post-surgery for NIC patients (Fig.-1a), or were drawn from eleven age and gender matched healthy donors (HD) obtained from the French blood donation center. Five bone marrow (BM) biopsies from orthopedic surgery patients were also included in this study. All specimens were obtained in accordance with the Declaration of Helsinki.

### Mass cytometry analysis

Whole blood samples were processed within 3 hours after sampling (Fig.-1b). A 42-dimensional mass cytometry panel was used. It included antibodies (Ab) either purchased ready tagged from Fluidigm or else purchased protein-free from the suppliers as indicated in Supplemental Table 3 and conjugated in house using MaxPar labelling kits from Fluidigm according to the manufacturer’s instructions.

A multi-step staining protocol was set up with 500μl of whole blood. Live surface staining was performed with metal-tagged Ab (m-Ab) targeting surface markers after Fc-blocking and Rhodium staining steps for live-dead cells discrimination as previously described (38, 39). Red blood cells were then lysed for 15min using BD PharmLyse solution (BD bioscience) and remaining leukocytes were permeabilized using Transcription Factor Buffer Set (BD bioscience) following manufacturer instructions. Permeabilized/fixed cells were then stained with m-Ab against intracellular molecules. Finally, cells were incubated over-night at +4°C with Iridium (250nM) in PBS containing 2% paraformaldehyde, for the detection of intact nucleated cells, and then directly frozen at −80°C. Once the collection of samples were completed, stained cells were thawed by batches including at least two samples from each group each time. Cells were washed once with PBS, then twice with deionized water and resuspended in deionized water containing standard normalization beads (Fluidigm) (40) at cell concentration adjusted to 5 × 10^5^ cells/ml. Cells were collected on a CyTOF Helios instrument located in the Cytometry platform (CyPS) of Sorbonne University. Acquired data were normalized with MATLAB-based normalization software (40), and were arcsinh transformed, prior to analysis using the Cytobank platform (41).

### Computationnal data analysis of all circulating cells

To identify the circulating immune populations, we used The Visualization of t-Distributed Stochastic Neighbor Embedding (viSNE implementation of t-SNE) (23) to automatically arrange cells according to their expression profile of 40 measured proteins (Supplemental Table 3) and to visualize all cells in a 2D map where position represents local phenotypic similarity. Next, expert gating on generated t-SNE axes was used to identify neutrophils and non-neutrophil cells.

### Computational data analysis of neutrophils

The neutrophils analysis started with a computational “discovery strategy” that aims to identify sepsis-specific subsets (Figure 1B), followed by a computational “validation strategy” (Figure 1C).

In the discovery strategy, the neutrophils from all patients were combined (18,800 neutrophils per sample from 67 samples) and embedded in a new set of t-SNE axes designated t-SNE1-2 and t-SNE2-2 according to per-cell expression of 7 key proteins (Supplemental Table 3). After dimensionality reduction, cells were clustered using Spanning-tree Progression Analysis of Density-normalized Events (SPADE) to group the neutrophils based on their high-dimensional expression profile into 55 nodes (24). These nodes were clustered according to log2-transformed cell abundance centered on the mean proportion of all samples’ nodes. Samples and mean-centered log2-transformed nodes cell proportion were depicted in a heatmap and arranged according to complete linkage hierarchical clustering (Supplemental Figure 3, Figure 2B). A second heatmap was generated to characterize cell nodes phenotype according to the mean expression of all panel markers (Supplemental Figure 4) and 7 chosen markers (CD11b, CD66b, CD10, CD16, CD64, CD123 and PD-L1) (Figure 2C). Markers were arranged according to complete linkage hierarchical clustering but nodes were ordered according to nodes order of Figure 2B heatmap.

In the validation strategy, Uniform Manifold Approximation and Projection (UMAP) (25) was used to create a single, common map of neutrophils across all samples (18,800 cells × 67 samples) using the 7 markers previously used (Supplemental Table3). Once a common two-dimensional representation of all samples was established, a FlowSOM clustering optimization was applied to determine optimal cluster number based on relative homogeneity of marker expression in clusters (42). The self-organized map generated in an unsupervised way with the optimized FlowSOM algorithm produced clusters that were stable and contained phenotypically homogenous cells (26). This strategy allowed the identification of 50 neutrophil clusters (Supplemental Figure 6C). The relative cell abundance was used to arrange samples’ groups according to complete linkage hierarchical clustering, as described for the first strategy (Supplemental Figure 6B). To phenotypically characterize these nodes, Marker Enrichment Modeling (MEM) (27, 28) was used. MEM quantified the enriched features of the 50 clusters identified by FlowSOM in a compact label of cell identity. Protein enrichment was reported on a +10 to −10 scale, where +10 indicates that the protein’s expression was especially enriched and −10 indicated that the protein’s expression was especially lacking from those cells, relative to the other neutrophil clusters (Supplemental Figure 6D).

### Computationnal data analysis of non-neutrophil cells

A t-SNE was run on the immune cells from the 67 samples of the study, excluding neutrophils, using 40 of the measured markers (Supplemental Table 3) and clustered into 200 nodes using SPADE (Supplemental Figure 1B). A heatmap was generated to characterize nodes phenotype according to the mean expression of the all the markers, where markers were arranged in columns and nodes in rows according to complete linkage hierarchical clustering (Supplemental Figure 8A). To identify an early sepsis-specific signature within these immune populations, we then compared the abundance of the 200 identified cell nodes of these immune populations between HD, NIC and S samples at day-1 using two t-test comparison of the log-2 transformed proportions (log2Fold > 1, *p*<0.01). The first test was done between HD and S at day-1 and the second between NIC and S at day-1then the nodes that were differentially abundant between HD and S at day-1 were comparated to the ones that were differentially abundant between NIC and S at day-1 (Supplemental Figure 8C).

### Flow cytometry validation panel

To validate new sepsis-specific neutrophils subsets a seven markers panel was designed for conventional florescent flow cytometry. 100μl blood from newly recruited sepsis patients was incubated with 30μl of fluorescent surface Ab mix (Supplemental Table 4) at room temperature for 20 minutes protected from light. Red blood cells were lysed and remaining cells fixed using BD Lysing solution (BD bioscience) for 15 minutes at room temperature. After one PBS wash, the samples were kept at +4°C and analyzed within 24h with BD LSRFortessa X20 flow cytometer.

### Activation and phagocytosis assay

To address neutrophils activation and phagocytic capacities we used pHrodo-labeled BioParticles and coated with *Staphylococcus aureus* (*S. aureus*) or Zymosan antigens (Invitrogen). pH-sensitive pHrodo dyes are useful for visualizing acidification process during phago-lysosome formation (29). Lyophilized (*S. aureus*) and Zymosan pHrodo BioParticles were reconstituted in uptake Buffer (20 mM HEPES in HBSS, pH 7.4) to a concentration of 1 mg/ml and 0,5 mg/ml, respectively. Each sample was aliquotted (100 μl/aliquot for blood or 2 × 10^5^ cells/aliquot in 100μl of RPMI containing 10%SVF for BM). Aliquots were incubated 1h protected from light, with either 20μl of *S. aureus* pHrodo BioParticles or 40μl of Zymosan pHrodo BioParticles at 37°C with 5% CO_2_ for positive controls (PC) or were maintained on ice for negative controls (NC). After the incubation, 10μl of fluorescent surface Ab mix (Supplemental Table 4) was added to each aliquot and incubated on ice for 20 minutes. Red blood cells were lysed and remaining cells fixed using BD Lysing solution (BD bioscience) for 15 minutes at room temperature. After one PBS wash, the samples were kept at +4°C and analyzed within 24h with BD LSRFortessa X20 flow cytometer.

### Statistical information

Numeric data are given as median ± SD with the exception of Fig. 7 data that are given as mean ± SD. Nonparametric two-tailed Mann-Whitney test with a significance threshold of alpha (a=0.05) was used to compare differences in cellular abundance of cell subsets between two patients’ groups in Fig. 2-5, 7A and in Supp. Fig. 5B and MFI ratios in Fig. 7D, 7E. Nonparametric two-tailed Wilcoxon signed-rank test with a significance threshold of alpha (a=0.05) was used to compare differences in cellular abundance of cell subsets from two matched patients samples in Fig. 4B, 5A, 5B. Correlation between two data sets in Supp. Fig. 7 was assessed using linear regression and Spearman correlation test. Statistical tests were performed using commercial statistics software GraphPad7.

## Author contributions

AMK, BGC, AB and CC designed the study. AMK and NG performed experimental work. BGC, KG and HV provided clinical samples, pathological diagnosis and patient clinical data. AMK compiled patient data. AMK and AC run samples in the mass cytometer. AMK, SMB, SG and JMI performed data analysis. AMK, JMI and CC developed the figures, and wrote the manuscript. CC provided financial support. All authors contributed in reviewing the manuscript.

## Acknowledgments

We thank Drs Nicolas Mongardon, Adrien Bouglé, Alice Blet, Pierre Mora, Nicolas Deye and Paul Delval from Assistance-Publique Hôpitaux de Paris, Paris, France, and Dr Delphine Sauce from Cimi-Paris for their help in samples collection.

## Funding

This work was supported by grants from Inserm, Sorbonne University, Fondation pour la recherche Médicale “Equipe labelisée” and from “Agence Nationale de la Recherche”, project CMOS (CX3CR1 expression on monocytes during sepsis) 2015 (ANR-EMMA-050). AMK was supported by post-doctoral fellowship both from the ANR and FRM.

## Competing interests

J.M.I. is a co-founder and a board member of Cytobank Inc. and received research support from Incyte Corp, Janssen, and Pharmacyclics.

## Notes

### Competing Interest Statement

Jonathan M. Irish is a co-founder and a board member of Cytobank Inc. and received research support from Incyte Corp, Janssen, and Pharmacyclics.

