## Supplemental figures and tables for "Two new immature and dysfunctional neutrophil cell subsets define a predictive signature of sepsis useable in clinical practice"

### Supplementary Materials:

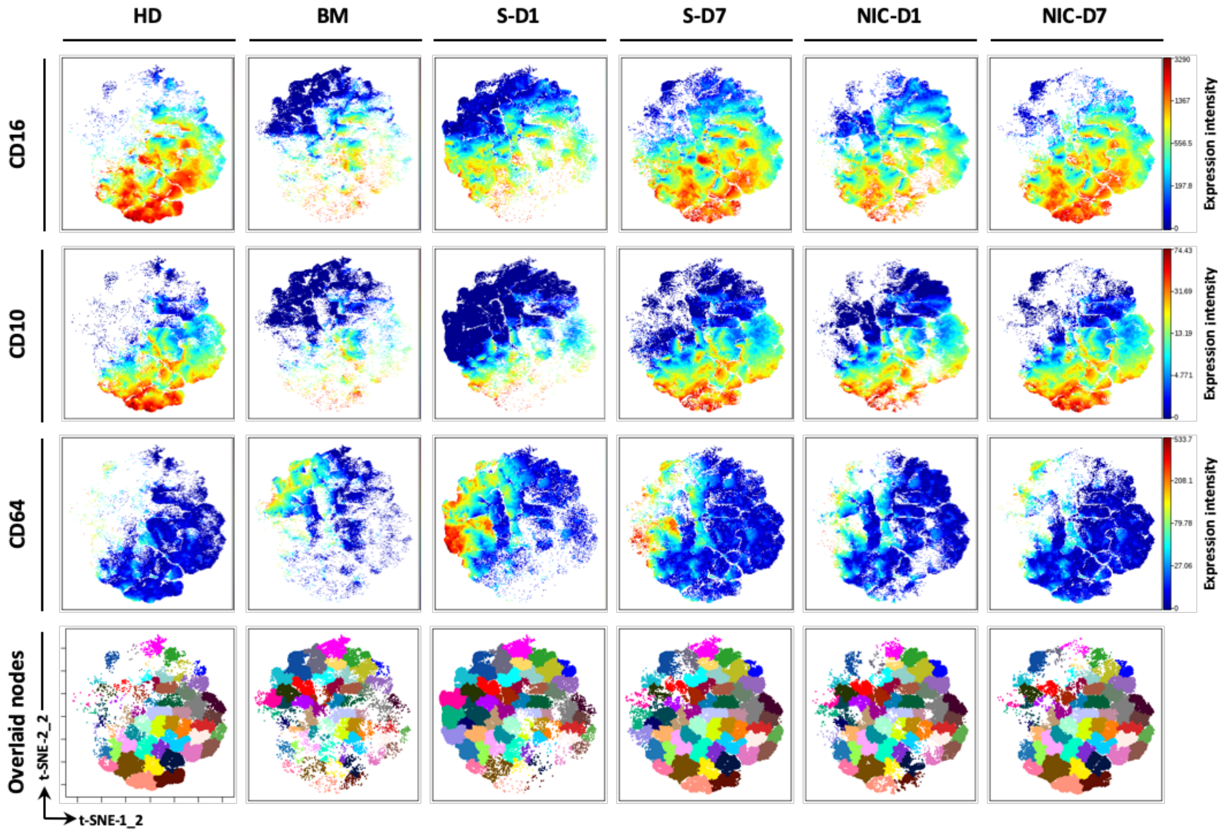


**Supplemental Figure 1. Neutrophils heterogeneity assessed by automatic clustering after dimensional reduction**. t-SNE analysis was performed on neutrophils from all samples with cells being organized along t-SNE-1-2 and t-SNE-2-2 according to per-cell expression of CD11b, CD66b, CD16, CD10, CD64 and CD123, PD-L1. Neutrophils' CD16, CD10, CD64 markers expression is presented on a rainbow heat scale in the t-SNE map of each group concatenated file. SPADE clustering was done to separate neutrophils subsets in 55 nodes. SPADE nodes of merged files of each group were overlaid on t-SNE map. t-SNE automatically arranged S patients' blood neutrophils on the upper left area of the map whereas the NIC patients' cells were in the central/upper right area. HD neutrophils were located on the lower right area of the map. Most of day-7 samples cells were gathered around HD area. BM cells were localized in the upper left quadrant in proximity to S day-1 cells.

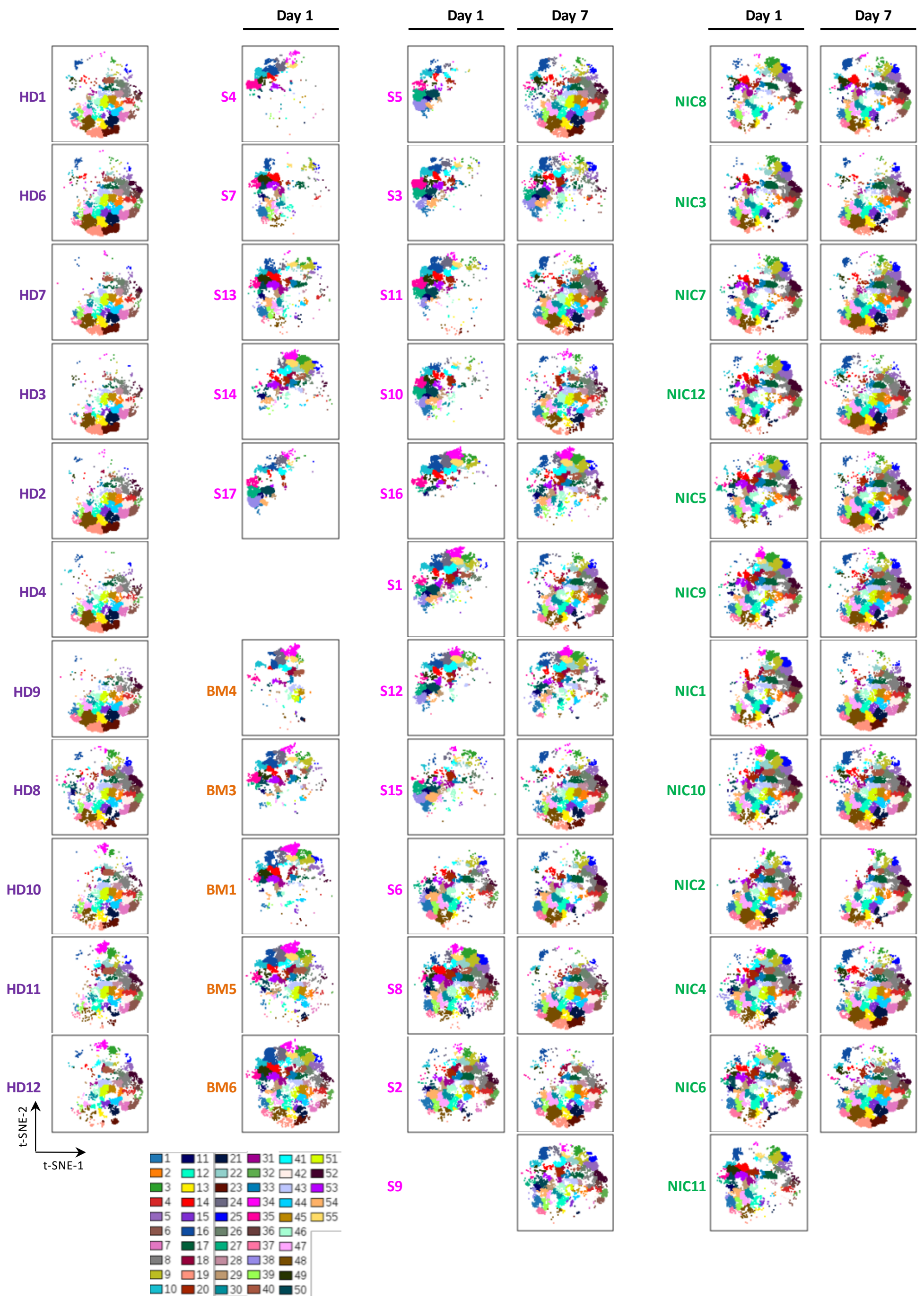


**Supplemental Figure 2. Neutrophils subsets evolution over time and between groups.** A second t-SNE analysis was performed on all samples neutrophils and cells were organized along a second couple of t-SNE axes (t-SNE-1-2 and t-SNE-2-2) according to per-cell expression of CD11b, CD66b, CD16, CD10, CD64 and CD123, PD-L1. SPADE clustering was done to separate neutrophils subsets in 55 nodes. The 55 SPADE nodes of each sample were overlaid on t-SNE map.

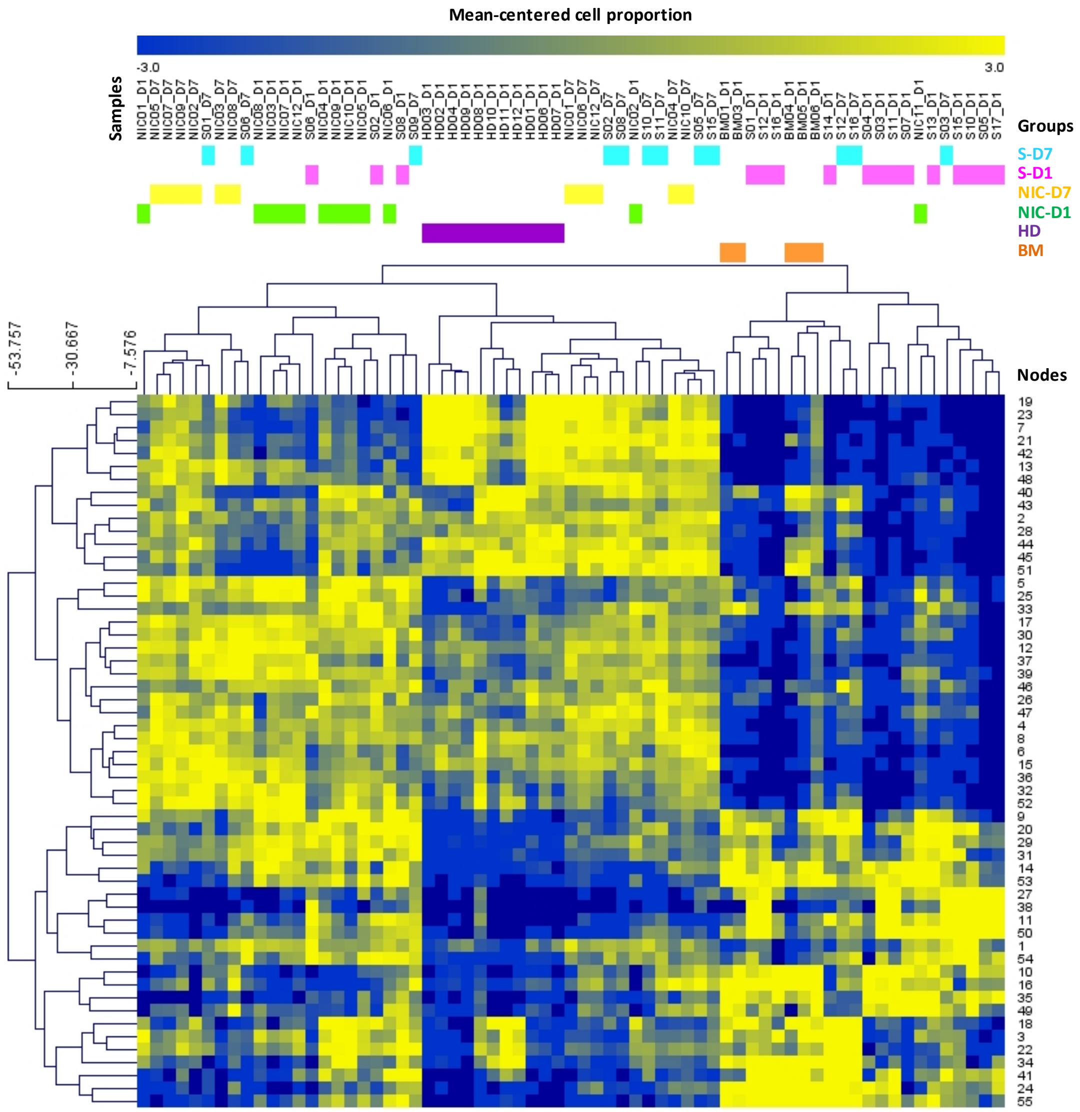


**Supplemental Figure 3. Identification of group-specific nodes with a heatmap analysis.** A heat map shows samples clustering (columns) according to nodes cell proportion log2-transformed and centered around the mean proportion of all samples' nodes (rows). Samples and mean-centered log2-transformed nodes cell proportion were arranged according to complete linkage hierarchical clustering. Heat intensity (from blue to yellow) reflects the mean-centered log2-transformed cell proportion of each sample's node.

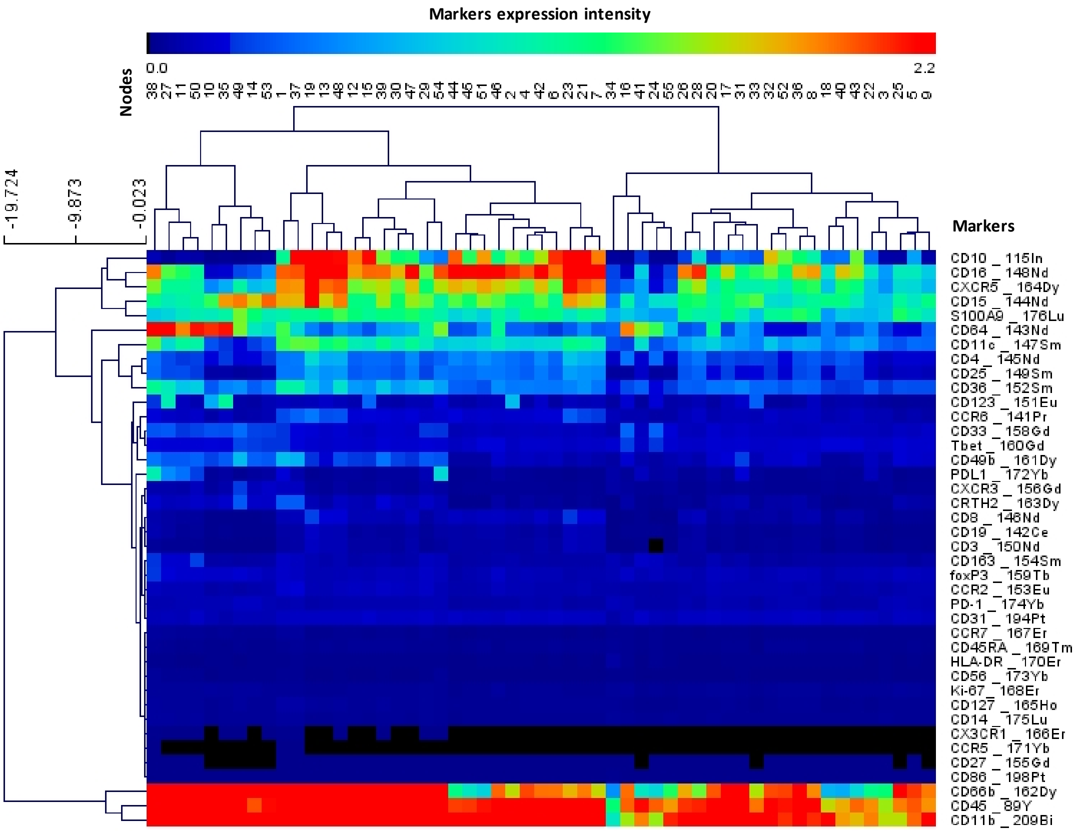


**Supplemental Figure 4. Characterizing cell nodes phenotype with a heat map analysis.** A heatmap shows characterization of cell nodes identified by SPADE (columns) according to mean expression of 40 markers (rows). Nodes and mean markers expression were arranged according to complete linkage hierarchical clustering. Heat intensity reflects the mean expression of each marker in each node.

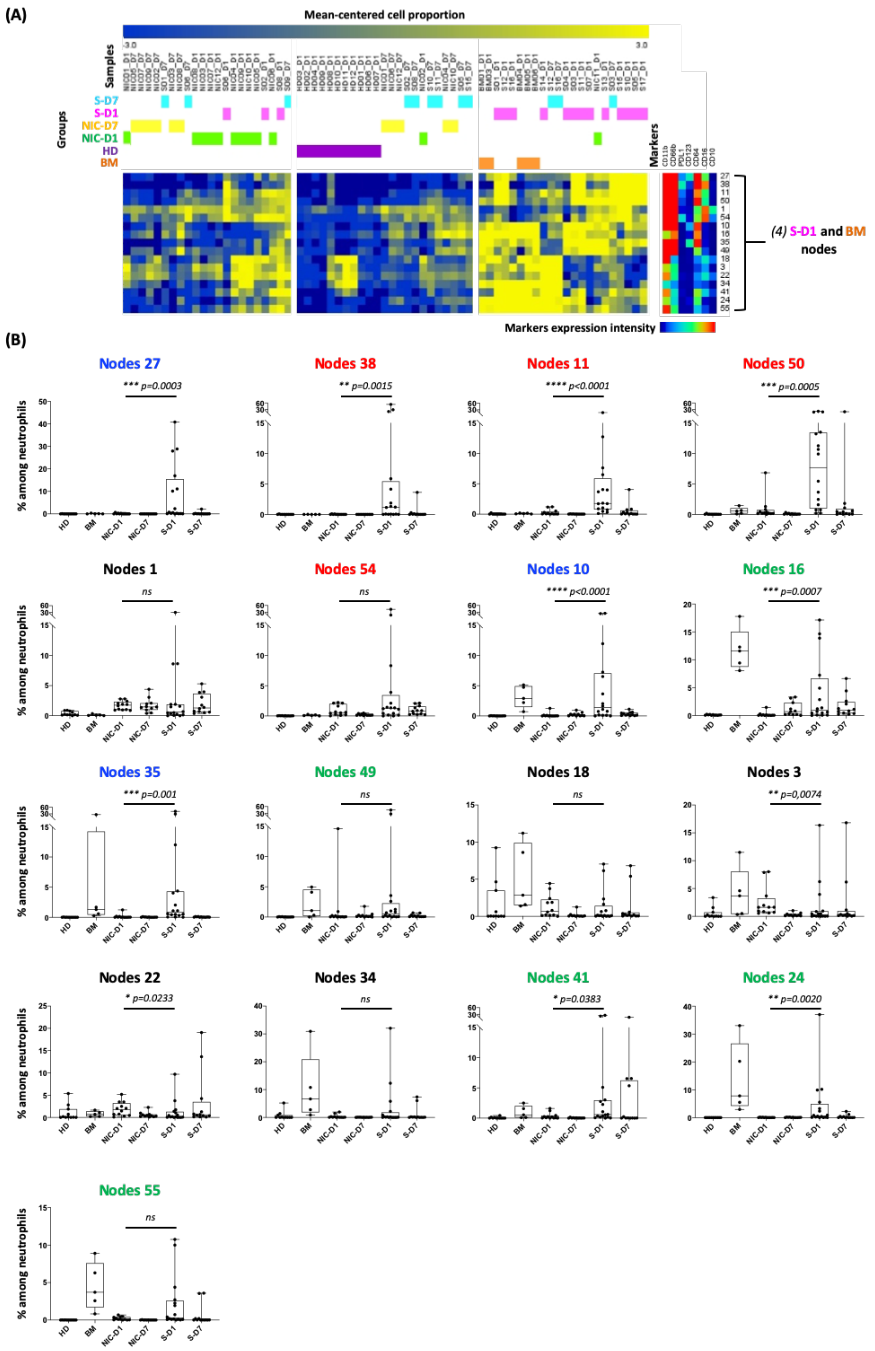


**Supplemental Figure 5. Neutrophils group (4) nodes proportion between.** Cell proportion of each of sepsis-Day-1 (S-D1) and bone marrow (BM)-enriched nodes, gathered in group (4) (**A**), was represented in (**B**) for each group. Statistics: Nonparametric two-tailed Mann-Whitney test was used to compare differences in cellular abundance of cell subsets between NIC-D1 and S-D1 (see the Methods section). Sample sizes: HD=12, BM=5, NIC-D1=12, NIC-D7=11, S-D1=16 and S-D7=12.

(a)

(b)

(c)

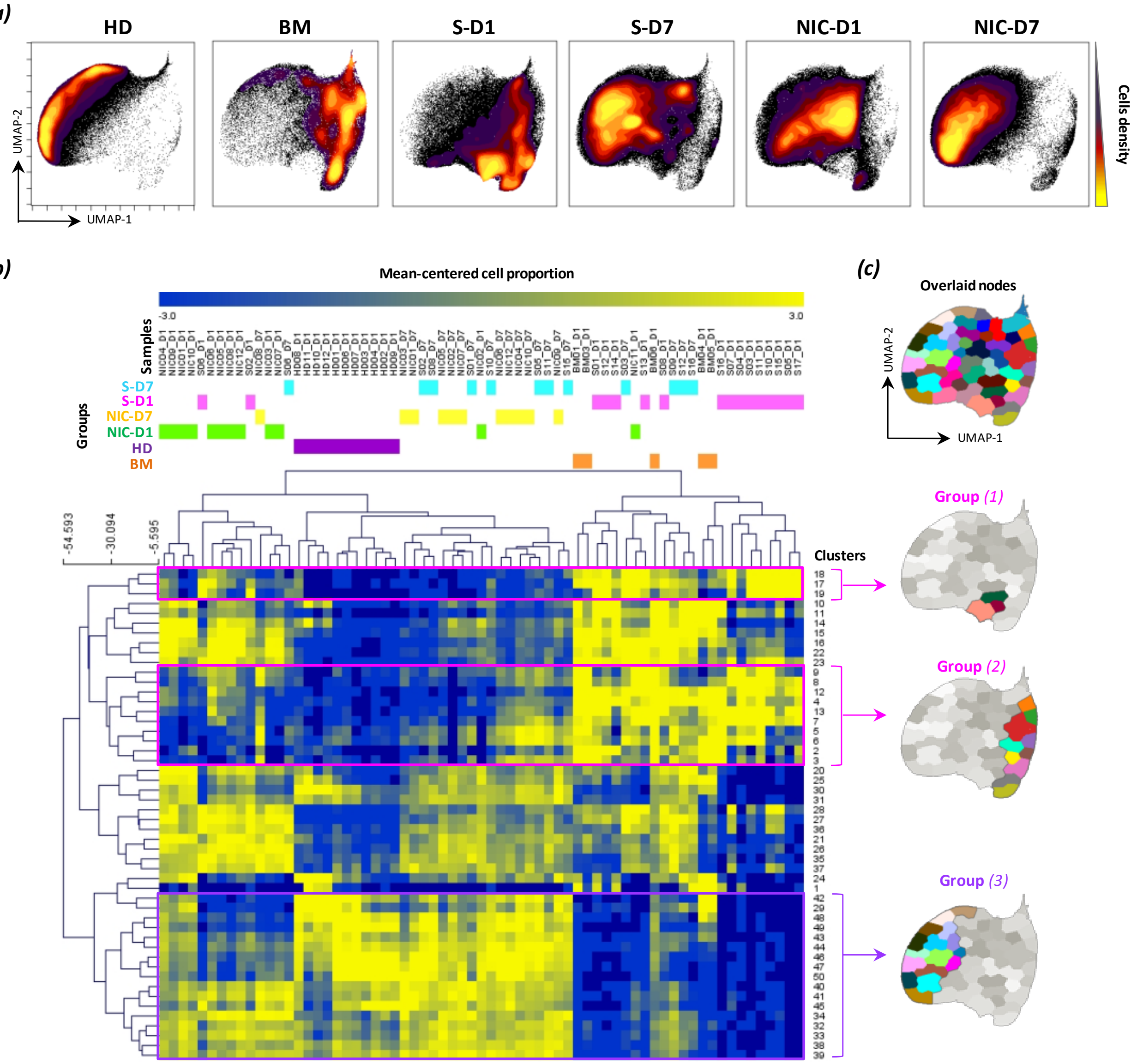


(d)

**Group *(1)* MEM Labels**

**Cluster 18:**
▲ CD64$^{+5}$ CD66b$^{+3}$ PD-L1$^{+1}$ CD123$^{+1}$
▼ CD10$^{-8}$ CD16$^{-5}$

**Cluster 17:**
▲ CD64$^{+4}$ CD66b$^{+2}$ PD-L1$^{+2}$
▼ CD10$^{-8}$ CD16$^{-2}$ CD11b$^{-2}$

**Cluster 19:**
▲ CD64$^{+6}$ CD66b$^{+2}$ PD-L1$^{+2}$
▼ CD10$^{-9}$

**Group *(2)* MEM Labels**

**Cluster 9:**
▲ CD64$^{+3}$ CD66b$^{+8}$ CD11b$^{+2}$
▼ CD10$^{-8}$ CD16$^{-7}$ CD11c$^{-3}$

**Cluster 8:**
▲ CD64$^{+3}$ CD66b$^{+6}$ CD11b$^{+1}$
▼ CD10$^{-8}$ CD16$^{-7}$ CD11c$^{-3}$

**Cluster 12:**
▲ CD64$^{+4}$
▼ CD10$^{-8}$ CD16$^{-4}$ CD66b$^{-2}$ CD11b$^{-2}$

**Cluster 4:**
▲ CD64$^{+4}$
▼ CD10$^{-8}$ CD16$^{-7}$ CD66b$^{-4}$ CD11b$^{-3}$

**Cluster 13:**
▲ CD64$^{+4}$ CD66b$^{+4}$
▼ CD10$^{-8}$ CD16$^{-7}$ CD11b$^{-3}$

**Cluster 7:**
▲ CD64$^{+4}$ CD66b$^{+3}$ CD123$^{+3}$
▼ CD10$^{-8}$ CD16$^{-7}$

**Cluster 5:**
▲ CD64$^{+5}$ CD66b$^{+3}$
▼ CD10$^{-8}$ CD16$^{-8}$ CD11c$^{-4}$ CD11b$^{-3}$

**Cluster 6:**
▲ CD64$^{+5}$ CD66b$^{+1}$ CD123$^{+1}$
▼ CD10$^{-8}$ CD16$^{-8}$

**Cluster 2:**
▲ CD64$^{+3}$
▼ CD10$^{-8}$ CD16$^{-8}$ CD66b$^{-4}$ CD11b$^{-6}$ CD11c$^{-4}$

**Cluster 3:**
▲ CD64$^{+3}$
▼ CD10$^{-8}$ CD16$^{-8}$ CD66b$^{-4}$ CD11b$^{-8}$ CD11c$^{-4}$

**Supplemental Figure 6. Validation of sepsis day 1-specific clusters by UMAP dimensional reduction, FlowSOM clustering and MEM labeling.** (**A**) UMAP analysis was performed on neutrophils from all samples and cells were organized along UMAP-1 and UMAP-2 axes according to per-cell expression of CD11b, CD66b, CD16, CD10, CD64 and CD123, PD-L1. Cell density for the concatenated file of each group is shown, on a black to yellow heat scale, for each group time-point. (**B**) FlowSOM clustering was then applied to separate neutrophils subsets into 50 clusters. A heat map shows samples clustering (columns) according to clusters cell proportion log2-transformed and centered around the mean proportion of all samples' nodes (rows). Samples and mean-centered log2-transformed nodes cell proportion were arranged according to complete linkage hierarchical clustering. Heat intensity (from blue to yellow) reflects the mean-centered log2-transformed cell proportion of each sample's node. (**C**) FlowSOM clusters of merged files of all samples and three groups of nodes were back-viewed on UMAP-1 / UMAP-2 map. (**D**) Marker Enrichment Modeling (MEM) was used to phenotypically characterize group (1) and group (2) clusters. Protein enrichment for each cluster was reported on a +10 to −10 scale, where +10 indicates that the protein's expression was especially enriched and −10 indicated that the protein's expression was excluded from those cells, relative to the other neutrophil clusters.

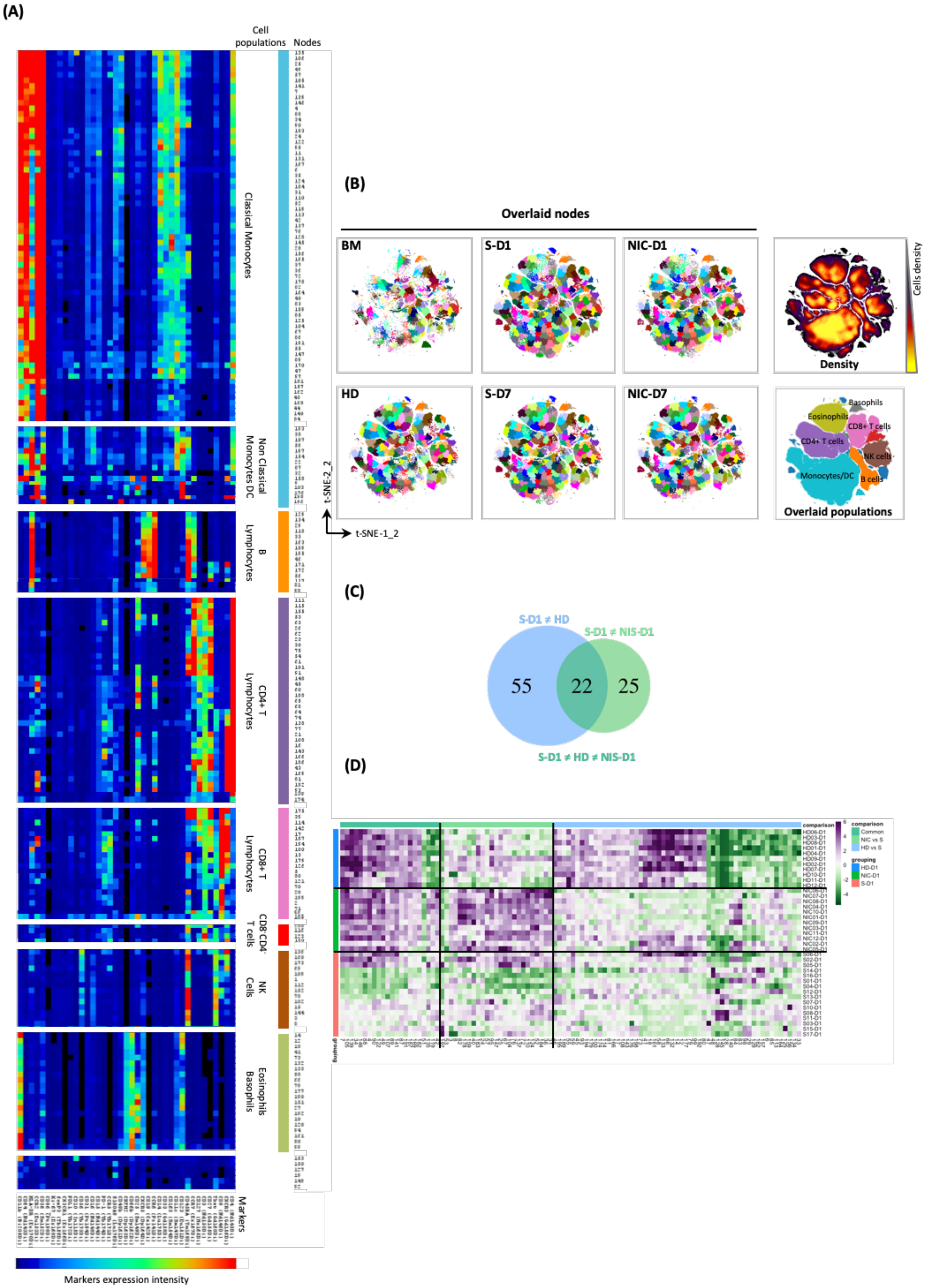
(A)
Cell populations
Nodes
Classical Monocytes
Non Classical Monocytes DC
B Lymphocytes
CD4+ T Lymphocytes
CD8+ T Lymphocytes
CD8-CD4- T cells
NK Cells
Eosinophils Basophils
Markers
Markers expression intensity
(B)
Overlaid nodes
BM
S-D1
NIC-D1
HD
S-D7
NIC-D7
Density
Cells density
Basophils
Eosinophils
CD8+ T cells
CD4+ T cells
NK cells
Monocytes/DC
B cells
Overlaid populations
t-SNE-2_2
t-SNE-1_2
(C)
S-D1 ≠ HD
S-D1 ≠ NIS-D1
55
22
25
S-D1 ≠ HD ≠ NIS-D1
(D)
comparison
Common
NIC vs S
HD vs S
grouping
HD-D1
NIC-D1
S-D1

**Supplemental Figure 7. Non-neutrophil cells computational analysis. (A)** t-SNE analysis was performed on all non-neturophil cells from all samples with cells being organized along the t-SNE-1-2 and t-SNE-2-2 axes according to per-cell expression of 40 markers. SPADE clustering was done to separate non-neutrophils immune cell subsets into 200 nodes. A heatmap shows characterization of cell nodes identified by SPADE (rows) according to mean expression of 40 markers (columns). Markers and nodes were arranged according to complete linkage hierarchical clustering. Heat intensity (from blue to red) reflects the mean expression of each marker for each node. The nodes were grouped according to the main immune subsets that were overlaid on the t-SNE map in **(B)**. **(B)** SPADE nodes of concatenated files for each group were overlaid on t-SNE plot. Cell density for the concatenated cohort is shown, on a black to yellow heat scale. Main immune subsets were manually gated and overlaid on the t-SNE map. **(C)** Venn-Diagram was used to present nodes that are differently abundant in S Day-1 group (SD1) when compared to HD but not to NIC Day-1 (NICD1) group (blue circle), and those that are differently abundant in SD1 when compared to NICD1 but not to HD (light green circle). The nodes that different from both HD and NICD1 are found in the intersection of the two circles (dark green). **(D)** A heat map shows samples clustering (rows) according to **(C)** nodes cell proportion log2-transformed and centered around the mean proportion of SD1 nodes (columns). Samples and mean-centered log2-transformed nodes cell proportion were arranged according to complete linkage hierarchical clustering. Heat intensity (from green to purple) reflects the mean-centered log2-transformed cell proportion of each sample's node.

| | CD66b | CD11b | CD64 | CD16 | CD10 |
|---|---|---|---|---|---|
| Neutrophils | ++ | ++ | - | +++ | ++ |
| Band cells | ++ | ++ | - | ++ | - |
| Metamyelocytes | ++ | ++ | ++ | + | - |
| Myelocytes | +++ | ++ | ++ | - | - |
| Promyelocytes | +++ | - | + | - | - |
| Myeloblastes | - | - | + | - | - |
| GMP | - | - | ++ | - | - |

**Supplemental Table1. Neutrophil progenitors' phenotype.** Modified from 16. GMP : granulocyte monocyte progenitor cell, +++ : strong expression, ++ : moderate expression, + : weak expression, - : no expression

| Parameters | Groups | | |
|---|---|---|---|
| | S | NIC | HD |
| Number of patients | 17 | 12 | 11 |
| Age at admission (yr) – no. ($Q_1$-$Q_3$) | 75 (70-83) | 75 (65-80) | 66 (63-70) |
| Male/female ratio | 10/7 | 8/4 | 8/3 |
| Admission category – Medical/Surgery | 8/9 | 0/12 | |
| Comorbidities (n) – 0/≥1 | 3/14 | | |
| SAPS II – Med ($Q_1$-$Q_3$) | 52 (37.5-64.5) | | |
| SOFA – Med ($Q_1$-$Q_3$) | 7.5 (3-10.5) | | |
| Site of infection | | | |
| Abdominal | 9 | | |
| Urinary | 2 | | |
| Pulmonary | 4 | | |
| Other | 2 | | |
| Type of infection | | | |
| Community acquired | 14 | | |
| Nosocomial | 3 | | |
| Microbiologically documented | | | |
| Bacilli Gram Negative | 11 | | |
| Cocci Gram Positive | 2 | | |
| Other | 4 | | |
| D7 non survivors, n(%) | 3 (18) | 1(8) | |
| D28 mortality, n(%) | 4 (23) | 1(8) | |
| Patient under MV n(%) | 13 (76) | 11 (92) | |
| Patient under vasopressors n(%) | 14 (82) | 5 (42) | |
| Immunologic parameters (CyTOF) - Med ($Q_1$-$Q_3$) | | | |
| mHLA-DR D1 | 150 (72-373)* | 327 (151-580)* | 708 (582-752) |
| Lymphocytes count (10*9/L) D1 | 0.3 (0.2-0.4)* | 0.3 (0.1-0.4)* | 0.8 (0.5-1.6) |
| Neutrophils count (10*9/L) D1 | 11.4 (5.5-15.3)* | 14.1 (9.9-17.4)* | 5.0 (3.4-5.8) |
| Neutrophils/Lymphocytes ratio D1 | 32.4 (18.9-65.4)* | 47.5 (29.1-94.2)* | 7.8 (3.7-8.4) |

**Supplemental Table2. Demographic, clinical, and immunologic data for included patients and healthy donors.** Blood samples were obtained from 17 sepsis patients, one day and seven days after their admission at the ICU. Continuous data and biological parameters are presented as median and IQR (quartile1 – quartile3). For clinical parameters, categorical data are presented as number of cases and percentages among the total population. SAPS II was calculated after admission. SOFA score was measured after 24h of ICU stay. mHLA-DR correspond to the median level of HLA-DR expressed on circulating monocytes assessed by mass cytometry. Lymphocytes and neutrophils counts were calculated using leukocytes counts obtained from the clinic for each individual at day-1and the proportion of each subset assessed by mass cytometry. Statistics: nonparametric two-tailed Mann-Whitney test was used to compare values between HD and S or NIC groups. *: $p<0.05$.

| | Isotope | Marker | Clone | Source | Markers<br>Live cells viSNE<br>Non-neutrophils viSNE | <br>Neutrophils viSNE<br>Neutrophils UMAP |
|---|---|---|---|---|---|---|
| 1 | 141Pr | CCR6 | G034E3 | Fluidigm | x | |
| 2 | 142Nd | CD19 | HIB19 | Fluidigm | x | |
| 3 | 143Nd | CD64 | 104D2 | Biolegend+ Fluidigm | x | x |
| 4 | 144Nd | CD15 | W6D3 | Fluidigm | x | |
| 5 | 145Nd | CD4 | RPA-T4 | Fluidigm | x | |
| 6 | 146Nd | CD8a | RPA-T8 | Fluidigm | x | |
| 7 | 147Sm | CD11c | Bu15 | Fluidigm | x | |
| 8 | 148Nd | CD16 | 3G8 | Fluidigm | x | x |
| 9 | 149Sm | CD25 | 2A3 | Fluidigm | x | |
| 10 | 150Nd | CD3 | UCHT1 | BD + Fluidigm | x | |
| 11 | 151Eu | CD123 | 6H6 | Fluidigm | x | x |
| 12 | 152Sm | CD36 | 5-271 | Fluidigm | x | |
| 13 | 153Eu | CCR2 | K036C2 | Fluidigm | x | |
| 14 | 154Sm | CD163 | GHI/61 | Fluidigm | | |
| 15 | 155Gd | CD27 | L128 | Fluidigm | x | |
| 16 | 156Gd | CXCR3 | G025H7 | Fluidigm | x | |
| 17 | 158Gd | CD33 | WM53 | Fluidigm | x | |
| 18 | 159Tb | Foxp3 | 259D/C7 | Fluidigm | x | |
| 19 | 160Gd | Tbet | 4B10 | Fluidigm | x | |
| 20 | 161Dy | CD49b | P1E6-C5 | Fluidigm | x | |
| 21 | 162Dy | CD66b | 80H3 | Fluidigm | x | x |
| 22 | 163Dy | CRTH2 | BM16 | Fluidigm | x | |
| 23 | 164Dy | CXCR5 | 51505 | Fluidigm | x | |
| 24 | 165Ho | CD127 | A019D5 | Fluidigm | x | |
| 25 | 166Er | CX3CR1 | 2A9-1 | Biolegend + Fluidigm | x | |
| 26 | 167Er | CCR7 | G043H7 | Fluidigm | x | |
| 27 | 168Er | Ki-67 | B56 | Fluidigm | x | |
| 28 | 169Tm | CD45RA | HI100 | Fluidigm | x | |
| 29 | 170Er | HLA-DR | L243 | Fluidigm | x | |
| 30 | 171Yb | CCR5 | NP-6G4 | Fluidigm | x | |
| 31 | 172Yb | PD-L1 | 29E.2A3 | Biolegend + Fluidigm | x | x |
| 32 | 173Yb | CD56 | B159 | Fluidigm | x | |
| 33 | 174Yb | PD-1 | EH12.2H7 | Fluidigm | x | |
| 34 | 175Lu | CD14 | M5E2 | Fluidigm | x | |
| 35 | 176Yb | S100A9 | 1H9 | Biolegend + Fluidigm | x | |
| 36 | 209Bi | CD11b | ICRF44 | Fluidigm | x | x |
| 37 | 89Y | CD45 | HI30 | Fluidigm | | |
| 38 | 194Pt | CD31 | WM59 | Biolegend + Fluidigm | x | |
| 39 | 198Pt | CD86 | IT2.2 | Biolegend + Fluidigm | x | |
| 40 | 115In | CD10 | HI10a | Biolegend + Fluidigm | x | x |
| 41 | 103Rh | Viability | | Fluidigm | | |
| 42 | 191-193Ir | DNA | | Fluidigm | | |

**Supplemental Table3. CyTOF panel and markers used for non-supervised analysis**

| Marker | Fluorochrom | clone | Source | Phagocytosis panel | Validation panel |
|---|---|---|---|---|---|
| CRTH2 | FITC | BM16 | Biolegend | | x |
| CD123 | PE | 6H6 | Biolegend | | x |
| BioParticles | pHrodo Red | | Invitrogen | x | |
| CD66b | AlexaF-700 | G10F5 | BD | x | |
| CD11b | APCH7 | ICRF44 | BD | x | |
| CD16 | BV421 | 3G8 | BD | x | |
| CD64 | BV605 | 10.1 | Biolegend | | x |
| PD-L1 | BV711 | 29E.2A3 | Biolegend | | x |
| CD15 | BV786 | W6D3 | BD | x | x |
| CD10 | BUV395 | HI10a | BD | x | x |
| CD14 | BUV737 | M5E2 | BD | | x |

**Supplemental Table4. flow cytometry panels**
